# A direct, MFRN-independent Fe(II) transfer pathway at mitochondria-lysosome contacts

**DOI:** 10.64898/2026.09.02.748755

**Authors:** Yubing Han, Xiaohai Hu, Xuelei Pang, Rongle Qiu, Mingwei Tang, Zhiyi Liu, Cuifang Kuang, Yu-Hui Zhang, Xu Liu

## Abstract

Mitochondrial iron homeostasis is fundamental to respiration and redox balance, and its dysregulation is implicated in neurodegeneration, cardiomyopathy, and metabolic diseases ^1, 2^. Although lysosomes harbor the major cellular iron reservoir ^3, 4^, the prevailing model holds that mitochondria acquire Fe(II) directly from the cytosolic labile iron pool (LIP)^5^ via MFRN transporters^6,7^. Here, we challenge the canonical view by identifying a direct, MFRN-independent Fe(II) transfer pathway at mitochondria-lysosome contacts (MLCs) ^8, 9^. This PPS39/TOMM22/SFXN1-coordinated pathway enables lysosome-to-mitochondria Fe(II) flux bypassing the cytosolic LIP. Using live-cell structured illumination microscopy (SIM), we visualize direct Fe(II) transfer specifically occurring at MLCs. Multiple lines of evidence confirm that PPS39 and TOMM22 stabilize MLCs, while SFXN1 serves as the core effector protein for this MLC-dependent Fe(II) transport. Notably, SFXN1 knockdown markedly reduces mitochondrial Fe(II) levels independent of its established serine transport function ^10^. This pathway reveals a major route for mitochondrial Fe(II) acquisition to support redox homeostasis^7^.

## Main

Free cytosolic Fe(II) acts as the predominant catalyst for Fenton chemistry, damaging nucleic acids, lipids, and proteins and imposing inherent metabolic costs upon the canonical LIP-dependent pathway of mitochondrial Fe(II) uptake^7, 11^. Membrane contact sites physically anchor paired organelles to mediate localized substrate shuttling, avoiding bulk cytosolic exposure and offering a feasible solution to mitigate cytotoxic oxidative risks.

Accumulating evidence demonstrates that mammalian mitochondria-lysosome contacts (MLCs) govern the trafficking of lipids, calcium, and multiple metabolites^12,13,14^. Given their role in bridging the major iron store to the primary iron consumer, MLCs may provide a platform for direct Fe(II) transfer between lysosomes and mitochondria^15,16^ (Fig. 1a). Despite this reasonable inference, the molecular basis of MLC-coupled Fe(II) translocation remains poorly understood^17^ in the classical LIP-MFRN-centric view. Based on our prior characterization of mitochondria-lysosome dynamic interactions^8^, here we integrate live-cell super-resolution imaging and genetic approaches to establish this lysosome-mitochondria Fe(II) transport pathway and the respective roles of PPS39^18^, TOMM22^19^, and SFXN1^20^ therein.

**Fig. 1.**
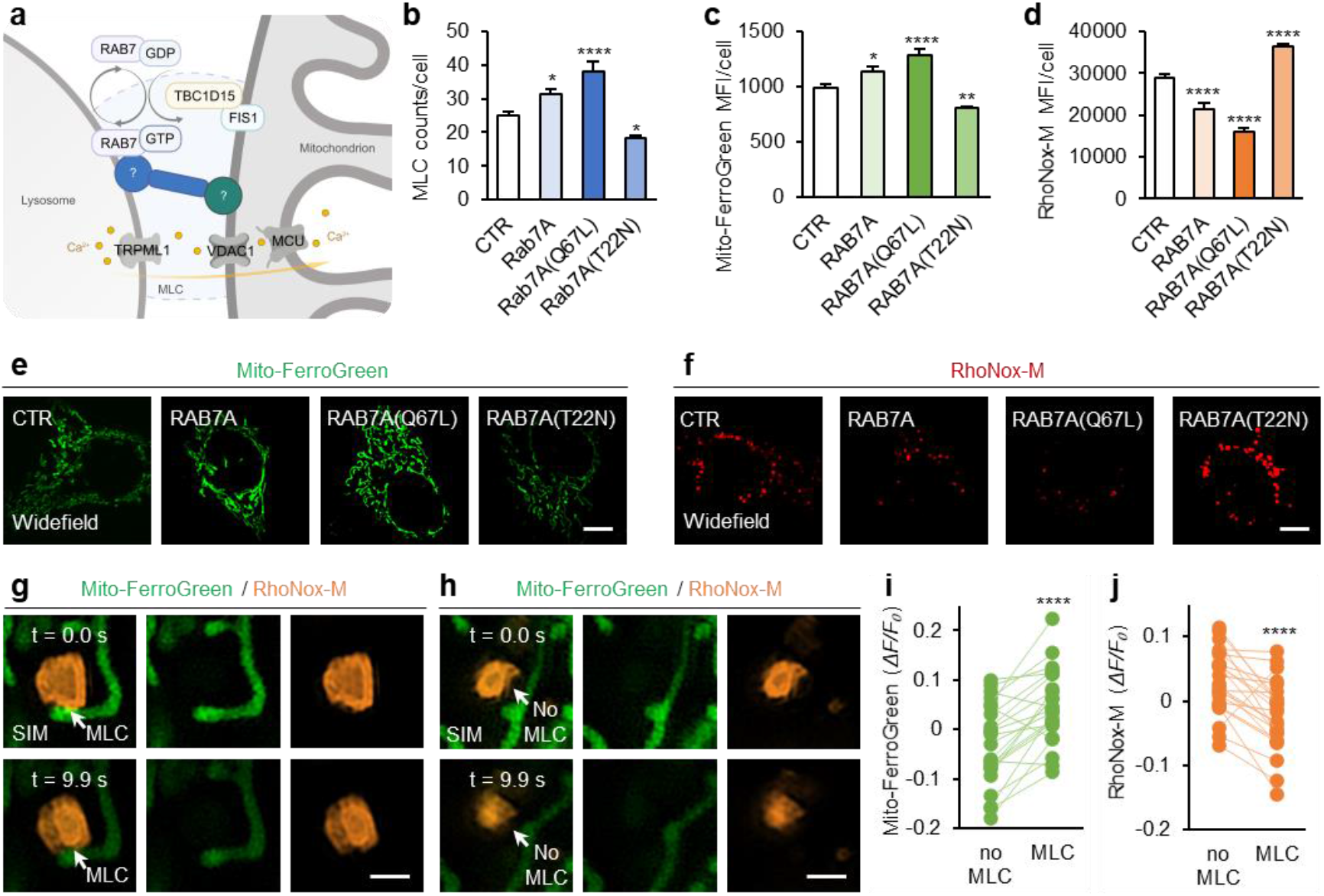
Mitochondria-lysosome contacts (MLCs) mediate Fe(II) redistribution between lysosomes and mitochondria. **a**, Established model of MLCs. RAB7 (regulated by FIS1/TBC1D15-mediated GTP hydrolysis) and associated tethering factors (unidentified components are indicated by question marks) promote MLC formation, creating a platform for inter-organellar communication, exemplified by Ca²⁺ transfer via the TRPML1-PDAC1-MCU pathway. **b**–**d**, Quantification of MLC puncta counts (**b**; from left to right: n =102, 106, 100, 99 cells), mitochondrial Fe(II) MFI (Mito-FerroGreen; **c**; from left to right: n = 91, 105, 105, 105 cells), and lysosomal Fe(II) MFI (RhoNox-M; **d**; from left to right: n = 94, 103, 105, 105 cells) under the indicated conditions. Statistical analysis was performed using one-way ANOPA with Tukey’s multiple comparisons test. **e**, **f**, Representative widefield images stained with Mito-FerroGreen (**e**) and RhoNox-M (**f**) as indicated. Scale bars, 10 μm. **g–j**, Time-lapse analysis of subcellular labile iron levels at MLC contacts and adjacent regions. MLCs were identified by SIM; Mito-FerroGreen and RhoNox-M signals were quantified from widefield images. Representative time-lapse SIM frames show MLC-contact regions (**g**) and paired no-MLC regions (**h**) within the same cell. Normalized fluorescence change (*ΔF/F₀*) values of Mito-FerroGreen (**i**) and RhoNox-M (**j**) signals were compared between MLC sites and paired no-MLC regions using two-tailed paired Student’s t-test (n= 23). Fluorescence signals were photobleaching-corrected using intensity from no-MLC areas. Scale bars, 1 μm. *P < 0.05, **P < 0.01, ***P < 0.001, ****P < 0.0001; ns, not significant.

### MLCs mediate direct lysosome-mitochondria Fe(II) transfer

To probe the contribution of MLC assembly to Fe(II) transfer, we transfected U2OS cells with wild-type (WT) RAB7, the constitutively active GTP-bound Q67L variant, or the GTPase-deficient T22N mutant^9, 21^. MLC abundance was elevated in cells expressing RAB7 (WT) and RAB7 (Q67L), while this contact structure decreased upon overexpression of RAB7 (T22N) variant (Fig. 1b, Supplementary Fig. 1). Consistent with this gradient of MLC levels, cells overexpressing RAB7 (WT) or RAB7 (Q67L) displayed elevated Mito-FerroGreen (mitochondrial Fe(II) probe) intensity and diminished RhoNox-M (lysosomal Fe(II) probe) fluorescence, whereas the opposite trend was observed in RAB7 (T22N)-overexpressing cells (Fig. 1c–f). Notably, RAB7 (WT) expression did not alter global cytosolic LIP levels (measured via FerroOrange) relative to control cells (Extended Data Fig. 1a,b), indicating that elevated mitochondrial Fe(II) is derived from lysosomal Fe(II) transport rather than overall LIP remodeling. Coordinated changes in LIP and mitochondrial Fe(II) were also observed in both RAB7 mutant lines (Extended Data Fig. 1a,b), possibly arising as a secondary consequence of disrupted lysosomal homeostasis driven by abnormal RAB7 hydrolytic activity.

We further verified direct Fe(II) transfer occurring at MLC structures using dual-color SIM imaging (Fig. 1g–j and Extended Data Fig. 1c–e). Quantitative analysis of Mito-FerroGreen and RhoNox-M fluorescence intensities was baseline-calibrated against photobleaching profiles measured from no-MLC subcellular compartments. Compared with no-MLC areas, MLC regions displayed increased Mito-FerroGreen normalized fluorescence change (*ΔF/F₀*), indicating a sustained Fe(II) supply to local mitochondria (Fig. 1i). This was accompanied by decreased lysosomal RhoNox-M fluorescence at MLC sites (Fig. 1j). Taken together with the preceding functional and molecular data, these observations support direct Fe(II) transfer across MLC interfaces.

### MLC stabilized by VPS39

The homotypic fusion and vacuole protein sorting (HOPS) ^22^ subunit PAM6/PPS39 binds RAB7 in yeast to mediate vacuole-mitochondria contact sites (vCLAMPs) ^23, 24, 25^. Yet whether PPS39 similarly supports MLC assembly in mammalian cells, and whether this assembly contributes to Fe(II) transfer, remained unexplored. Here we employed triple-color live-cell SIM to explore the functional role of PPS39 in regulating MLC biogenesis. Live-cell imaging showed that mCherry-PPS39 colocalized with lysosomal markers and supported stable, dynamic lysosome-mitochondrial tethering, where mitochondrial fission ^26^ and nascent mitochondrial branches^27^ were detected (Fig. 2a,b and Extended Data Fig. 2a–g).

**Fig. 2.**
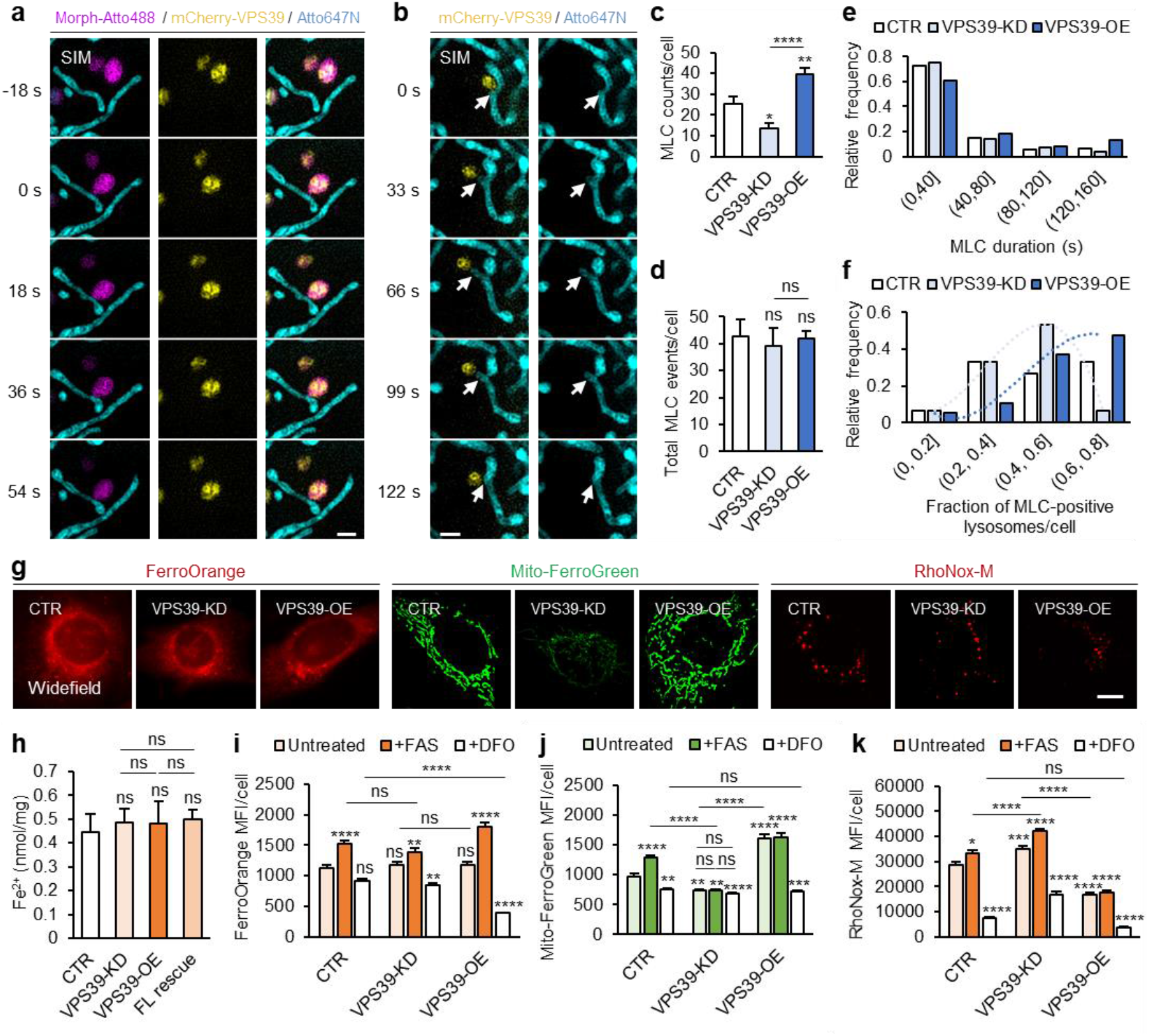
VPS39 localizes to MLCs and regulates direct Fe(II) transfer. **a**, **b**, Representative time-lapse structured illumination microscopy (SIM) images of stable MLCs in live cells stained with Atto647N (cyan; mitochondria), Morph-Atto488 (magenta; lysosomes), and mCherry-PPS39 (yellow). Scale bars, 1 μm. **c–f**, Quantification of MLC dynamics from time-lapse SIM images, showing the number of MLC contacts per cell quantified at the first frame (**c**; from left to right: n = 12, 12, 16 cells), total number of MLC events per cell accumulated across the full imaging duration (**d**; from left to right: n = 12, 12, 16 cells), relative frequency distribution of MLC contact duration (**e**; from left to right: n = 514, 468, 667 events), and or relative frequency distribution of MLC-positive lysosomes fraction per cell (**f**; from left to right: n = 12, 12, 16 cells). Overlaid third-order polynomial fit curves for PPS39-KD (light-blue) and PPS39-OE (dark-blue) revealing distinct distributions. **g–k**, Representative widefield images (**g**) and quantification of total iron (**h**; from left to right: n = 12, 12, 9, 9 cells; scale bar, 10 μm), LIP (FerroOrange; **i**; from left to right: n = 115, 105, 105, 109, 106, 105, 105, 105, 103 cells), mitochondrial iron (Mito-FerroGreen; **j**; from left to right: n = 102, 112, 109, 110, 107, 105, 106, 105, 105 cells), and lysosomal iron (RhoNox-M; **k**; from left to right: n = 110, 111, 105, 123, 106, 104, 105, 105, 105 cells) under the indicated PPS39 levels. Statistical analysis was performed using one-way ANOPA (**c**, **d**, **h**) and two-way ANOPA (**i–k**; details provided in Supplementary Tables 1–3) with Tukey’s multiple comparisons test. *P < 0.05, **P < 0.01, ***P < 0.001, ****P < 0.0001; ns, not significant.

To quantify PPS39’s regulatory role in MLC assembly, we established PPS39 knockdown (KD), overexpression (OE), and rescue cell lines (Extended Data Fig. 2h,i and 3), and performed time-lapse SIM imaging. Static quantification using the first frame of image sequences revealed that the number of MLCs per cell decreased upon PPS39 depletion, with re-expression restoring the defect (Fig. 2c and Extended Data Fig. 3). However, cumulative total MLC events over the entire imaging window were comparable across groups (Fig. 2d), indicating that PPS39 does not affect MLC initiation. By contrast, kinetic analysis from the same recordings showed that MLC duration was shortened in KD cells and prolonged in OE cells (Fig. 2e), accompanied by a corresponding shift in the distribution of MLC-positive lysosomes (Fig. 2f). Together, these data suggest that the static MLC abundance changes reflect altered MLC duration rather than initiation, consistent with the function of PPS39 as a tethering factor.

### VPS39 regulates MLC-dependent Fe(II) transfer

Building on the above findings, we further investigated whether PPS39 regulates Fe(II) transfer in the context of MLC function. Total intracellular Fe(II) and cytosolic LIP were comparable regardless of PPS39 levels (Fig. 2g–i). PPS39 KD reduced mitochondrial Fe(II) by 24% and increased lysosomal Fe(II) by 22%; both changes were reversed by PPS39 OE and rescued by PPS39 re-expression (Fig. 2g,j,k and Extended Data Fig. 4). The consistency between Fe(II) redistribution and MLC numbers indicates that PPS39 regulates lysosome-mitochondria Fe(II) transfer through MLC formation, independent of the cytosolic LIP pool.

We next performed pharmacological assays using ferrous ammonium sulfate (FAS) to enrich cytosolic LIP and deferoxamine (DFO) to chelate lysosomal labile Fe(III)^28^ (Fig. 2i–k and Extended Data Fig. 4 and Supplementary Tables 1–3). In control cells, FAS treatment increased both lysosomal and mitochondrial Fe(II), whereas DFO had the opposite effect. Upon PPS39 KD, however, FAS and DFO still altered lysosomal and LIP Fe(II) but failed to change mitochondrial Fe(II), indicating that loss of PPS39 decouples the mitochondrial Fe(II) pool from its upstream supply (Supplementary Note 1 for detailed discussion). Collectively, these data indicate that PPS39 regulates Fe(II) transfer between mitochondria and lysosomes via an MLC-dependent mechanism.

### Physical interaction and spatial proximity of VPS39, TOMM22, and SFXN1

To dissect the molecular mechanism underlying MLC-dependent lysosome-to-mitochondria Fe(II) transfer, we performed bioinformatic analysis including Gene Ontology Cellular Component (GO-CC) annotation and protein scoring criteria to screen candidate outer mitochondrial membrane (OMM) proteins (Extended Data Fig. 5a,b and Supplementary Note 2). Forward and reciprocal Co-IP assays in HEK293T cells confirmed a specific interaction between PPS39 and TOMM22 (Fig. 3a,b and Extended Data Fig. 5c,d). Given that TOMM22 functions as a OMM protein-conducting channel rather than an ion transporter, we intersected a library of PPS39 interactors with a predicted iron transporter dataset and identified the inner mitochondrial membrane (IMM) protein SFXN1 as a downstream candidate (Extended Data Fig. 5e,f and Supplementary Note 2). Biochemical validation confirmed that SFXN1 interacts with both PPS39 and TOMM22 (Fig. 3c–f).

**Fig. 3.**
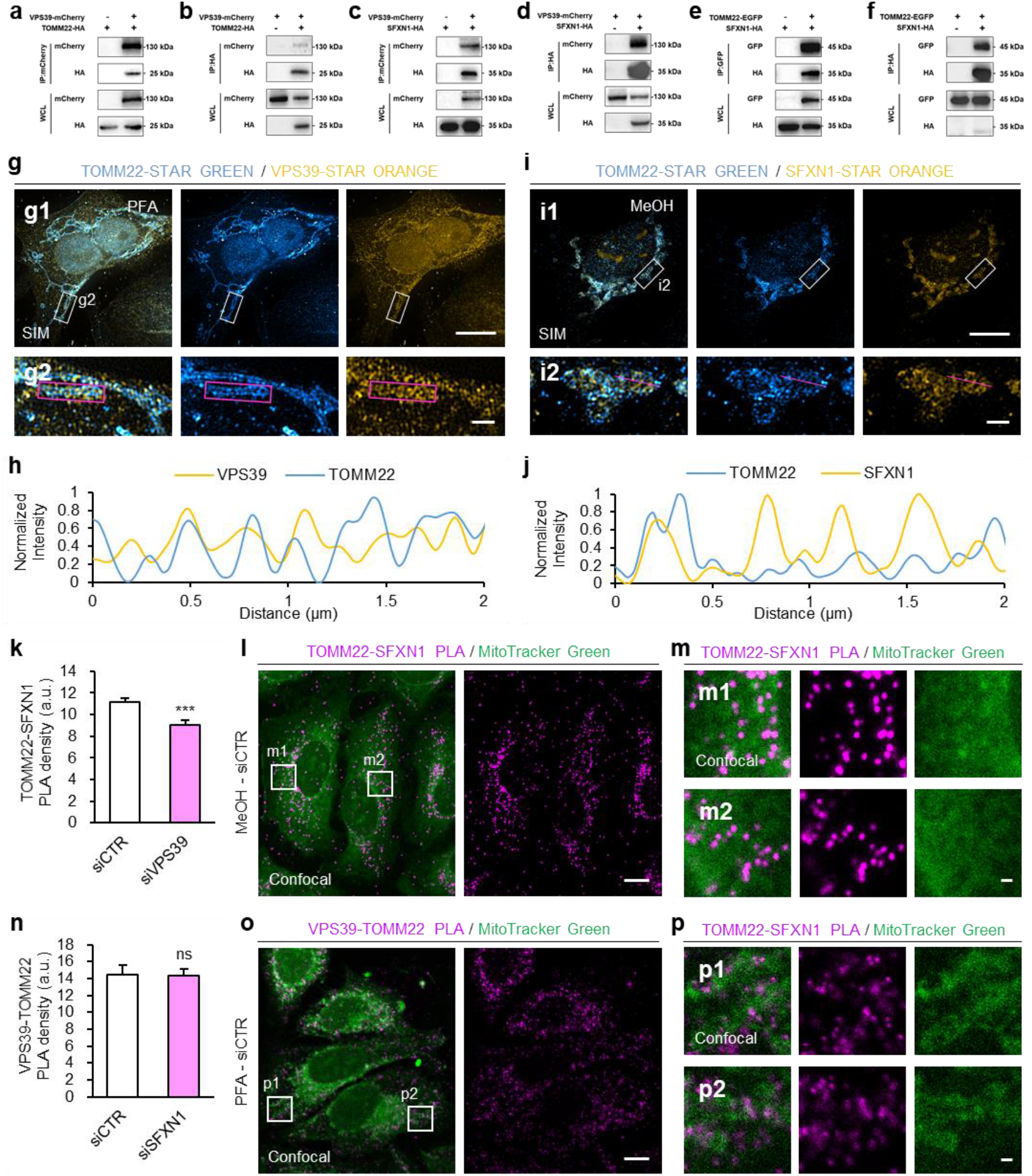
Interactions between VPS39, TOMM22 and SFXN1. **a–f**, Pairwise reciprocal co-immunoprecipitation (co-IP) assays of PPS39, TOMM22, and SFXN. **g–j**, Dual-color SIM immunofluorescence images of PPS39-TOMM22 (**g**; fixed with 4% paraformaldehyde) and TOMM22-SFXN1 (**i**; fixed in ice-cold methanol) complexes, with corresponding magenta intensity profiles shown in (**h**) and (**j**). Wide-field overviews (**g1**, **i1**; scale bars, 10 μm) and magnified views of the boxed regions (**g2**, **i2**; scale bars, 1 μm) are shown for each. SFXN1 exhibits a periodic distribution pattern that aligns with the organization of mitochondrial cristae. **k–p**, Proximity ligation assay (PLA) analysis of TOMM22-SFXN1 and PPS39-TOMM22 interactions. Quantification of PLA puncta abundance for PPS39-TOMM22 (**k**) and TOMM22-SFXN1 (**n**). Corresponding representative PLA images (**l**, **o**; scale bars, 10 μm) and the magnified views of the boxed regions (**m**, **p**; scale bars, 1 μm) from **l** and **o**. Statistical analysis was performed using two-tailed unpaired Student’s t-test. *P < 0.05, **P < 0.01, ***P < 0.001, ****P < 0.0001; ns, not significant.

We next performed dual-color SIM imaging to characterize the subcellular distribution of these three proteins. While PPS39 yields diffuse cytosolic signals and ring-shaped puncta morphology resembling lysosomal compartments, strikingly, both endogenous PPS39 and GFP-fused PPS39 produce faint yet reproducible signals outlining the TOMM22-related mitochondrial network (Fig. 3g,h and Extended Data Fig. 6a–e and Supplementary Note 3). Consistent with established knowledge^10,29^, OMM-associated TOMM22 and IMM-associated SFXN1 localize to distinct mitochondrial subdomains (Fig. 3i,j and Extended Data Fig. 6f).

### VPS39-TOMM22 support MLCs for Fe(II) transfer

We further interrogated whether the close spatial proximity resolved by super-resolution SIM facilitates complex formation among PPS39, TOMM22 and SFXN1. Co-IP assays in HEK293T cells revealed that siRNA-mediated silencing of either PPS39 or TOMM22 impairs the physical interactions between the retained protein and SFXN1, whereas SFXN1 depletion leaves PPS39-TOMM22 binding unaltered (Extended Data Fig. 7a–c). Consistently, cellular proximity ligation assay (PLA) quantification in U2OS cells reproduced these biochemical trends. PPS39 depletion lowered the abundance of TOMM22-SFXN1 PLA puncta (Fig. 3k–m and Extended Data Fig. 7d–f), while SFXN1 knockdown caused no obvious change to PPS39-TOMM22 PLA foci adjacent to MitoTracker-labelled structures (Fig. 3n–p and Extended Data Fig. 7g–i). Collectively, PPS39 and TOMM22 form a stable binary complex independent of SFXN1.

In consistence with PPS-depletion patterns, TOMM22 silencing reduced MLC abundance and impaired lysosome-mitochondria Fe(II) transfer, and both defects were fully rescued by TOMM22 re-expression (Extended Data Fig. 8), indicating that TOMM22 supports MLC abundance to facilitate inter-organelle Fe(II) transfer.

### Regulation of MLC-dependent Fe(II) transfer by SFXN1

In contrast, SFXN1 knockdown suppressed lysosome-mitochondria Fe(II) transfer (mitochondrial Fe(II) decreased by 57.8% relative to control groups), whereas MLC counts showed no statistically significant changes (Fig. 4a–e). As a documented mitochondrial serine permease, SFXN1 may exert extra regulatory control over matrix labile iron pools^10^. However, two lines of evidence from gradient L-serine supplementation in wild-type cells (Extended Data Fig. 9) functionally decouple these two activities. First, if altered iron homeostasis upon SFXN1 silencing merely represented a secondary consequence of impaired serine import, serine shortage would hinder downstream heme and Fe-S biogenesis and cause mitochondrial iron retention^30,31^, rather than the mitochondrial Fe(II) depletion we observed (Fig. 4c). Second, excess serine treatment only elevated cytoplasmic LIP without altering lysosomal Fe(II) levels (Extended Data Fig. 9d–f), which presents a distinct iron partitioning signature compared with SFXN1 knockdown. Collectively, these results support functional decoupling between SFXN1-mediated Fe(II) transport and serine translocation.

**Fig. 4.**
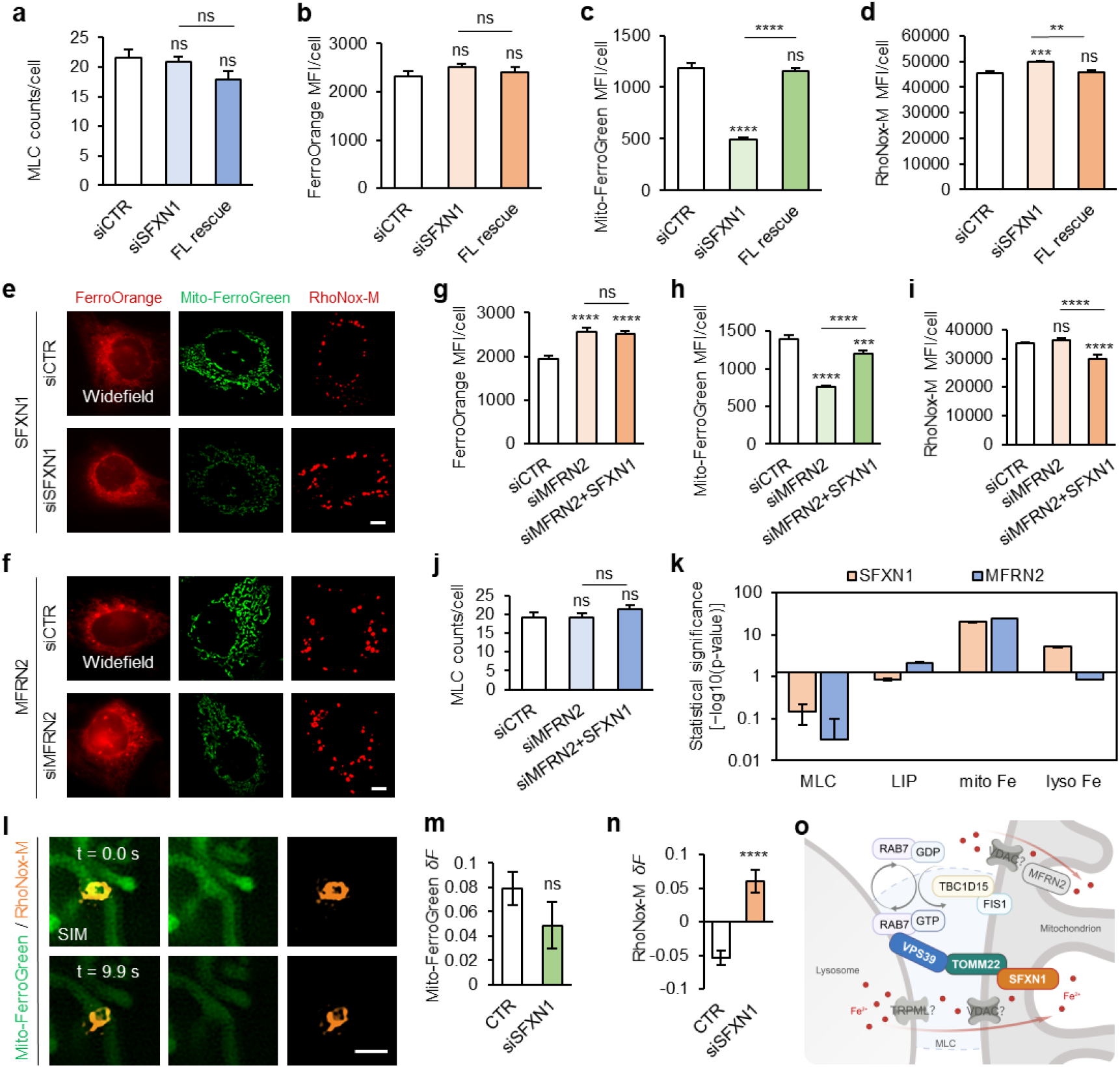
SFXN1 regulates MFRN2-independent Fe(II) transfer at MLCs. **a–d**, Quantification of MLC counts (**a**; from left to right: n = 105, 104, 98 cells), Fe(II) levels in the LIP (**b**; from left to right: n = 99, 99, 105 cells), mitochondria (**c**; from left to right: n = 109, 106, 105 cells), and lysosomes (**d**; from left to right: n = 105, 97, 105 cells) upon siRNA-mediated knockdown of SFXN1 (siSFXN1) and other indicated conditions. Statistical analysis was performed using one-way ANOPA with Tukey’s multiple comparisons test. **e**, **f**, Representative images of LIP, lysosomal iron, and mitochondrial Fe(II) upon siRNA-mediated knockdown of SFXN1 (**e**) and MFRN2 (**f**). Scale bar, 10 μm. **g–j**, Quantification of Fe(II) levels in the LIP (**g**; from left to right: n = 100, 104, 104 cells), mitochondria (h; from left to right: n = 101, 105, 105 cells), lysosomes (**i**; from left to right: n = 105, 105, 105 cells), and MLC counts (**j**; from left to right: n = 101, 99, 97 cells) upon siRNA-mediated knockdown of MFRN2 (siMFRN2) and other indicated conditions. Statistical analysis was performed using one-way ANOPA with Tukey’s multiple comparisons test. **k**, Divergent statistical significance patterns of organellar iron pools between SFXN1 and MFRN2 knockdown. The horizontal axis is set at Y = −log_10_(0.05) = 1.3, representing the significance threshold. Columns extending upward indicate statistical significance (p < 0.05); columns extending downward indicate non-significance (p > 0.05). Y-axis shows the original −log_10_(p) values. Data are derived from n ≈ 100 independent replicates per group; error bars represent 95% confidence intervals. MLC, MLC counts; LIP, cytosol labile iron pool (FerroOrange); Mito Fe, mitochondrial Fe(II) (Mito-FerroGreen); Lyso Fe, lysosomal Fe(II) (RhoNox-M). **l–n**, Time-lapse analysis of subcellular labile iron levels at MLC contacts and adjacent regions upon siSFXN1. MLCs were identified by SIM; Mito-FerroGreen and RhoNox-M signals were quantified from widefield images. Representative time-lapse frames show MLC-contact regions (**l**). Net difference in normalized fluorescence change (*ΔF/F₀*) for Mito-FerroGreen (**m**) and RhoNox-M (**n**). *δF* = *ΔF/F_0_*(MLC) − *ΔF/F_0_*(no MLC), calculated from MLC site and no-MLC ROIs within the same individual cell. *δF* values were then compared between CTR (n= 23) and siSFXN1 (n= 20) cells using two-tailed unpaired Student’s t-test. Scale bars, 1 μm. **o**, Schematic illustration of Fe(II) transport mediated by PPS39, TOMM22 and SFXN1 at MLCs. *P < 0.05, **P < 0.01, ***P < 0.001, ****P < 0.0001; ns, not significant. Data are pooled from three independent biological replicates.

To further verify the functional role of SFXN1 in lysosome-mitochondria Fe(II) transfer, we performed individual and combined knockdown of MFRN1 (mitoferrin 1) and MFRN2 (mitoferrin 2) for comparative analysis (Fig. 4f–k and Extended Data Fig. 10a–j). MFRN2 is the dominant mitochondrial Fe(II) transporter under steady state in non-erythroid cells^6^. Consistent with previous reports ^32^, knockdown of MFRN1, MFRN2, or combined MFRN1/2 all led to reduced mitochondrial Fe(II) and elevated cellular LIP Fe(II) relative to control cells, while no significant changes were observed in the number of MLCs or lysosomal Fe(II) levels (Fig. 4f–k and Extended Data Fig. 10a–d). Notably, overexpression of SFXN1 restored mitochondrial Fe(II) levels in all three MFRN-deficient backgrounds, accompanied by reduced lysosomal Fe(II) content, whereas cellular LIP levels and MLC counts remained unchanged (Fig. 4g–j and Extended Data Fig. 10a–d). Comparative analysis of SFXN1 and MFRN2 knockdown showed that while both suppressed mitochondrial Fe(II) levels, they triggered distinct Fe(II) redistribution, with SFXN1 disrupting lysosomal Fe(II) homeostasis and MFRN2 altering only the cellular LIP (Fig. 4k). These data indicate that lysosome-to-mitochondria Fe(II) transfer is specifically mediated by SFXN1 and operates independently of both MFRN1 and MFRN2.

We next visualized real-time Fe(II) dynamics at individual MLCs in SFXN1-knockdown cells via live-cell time-lapse SIM (Fig. 4l–n and Extended Data Fig. 10k–o). *δF* represents the net difference in normalized fluorescence change (*ΔF/F₀*) measured between MLC and no-MLC regions, where *δF* = *ΔF/F_0_* (MLC) - *ΔF/F_0_* (no MLC). Mito-FerroGreen *δF* exhibited a moderate downward trend in SFXN1-silenced cells (Fig. 4m). This change did not reach statistical significance, likely owing to inter-mitochondrial heterogeneity in iron and SFXN1 levels. RhoNox-M *δF* shifted from negative values in control cells to positive values in SFXN1-silenced cells at MLC sites (Fig. 4n), suggesting that lysosome-derived iron at MLC sites fails to fully transfer into the mitochondrial matrix and instead accumulates within lysosomes at these contact sites. Together with the finding that SFXN1 knockdown does not affect MLC abundance, these results indicate that SFXN1 is required for efficient lysosome-to-mitochondria Fe(II) redistribution at intact MLCs (Fig. 4o).

## Discussion

Our findings establish a previously unrecognized MLC-dependent route for Fe(II) transfer from lysosomes to mitochondria that operates independently of both the cytosolic LIP and the MFRN transporters. This pathway is molecularly and functionally distinct from the BDH2-mediated Fe(II) transfer recently reported at MLCs in melanoma cells, in which 2,5-DHBA-bound Fe(II) enters mitochondria via MFRN1^17^.

SFXN1 loss alone is not associated with general mitochondrial dysfunction^10^, and in this pathway, the mitochondrial Fe(II) level regulated by SFXN1 is independent of serine transport and MFRN-mediated uptake, pointing to SFXN1 as the core effector for lysosome-to-mitochondria Fe(II) transfer. Notably, relative to their respective control groups, SFXN1 depletion triggered a larger fractional reduction in mitochondrial Fe(II) than MFRN2 knockdown (Fig. 4c,h). Since these two perturbations involve separate upstream Fe(II) reservoirs and cannot guarantee equivalent knockdown efficiencies, we refrain from direct quantitative comparison between the two routes. Nevertheless, this observation identifies the MLC-dependent pathway as a major, if not predominant, route for mitochondrial Fe(II) acquisition in this cellular setting.

By bypassing the cytosolic LIP, this direct lysosome-to-mitochondria Fe(II) transfer may limit the risk of Fenton chemistry and associated oxidative damage^11^. Collectively, our study identifies promising molecular targets for a spectrum of disorders including neurodegeneration, cardiomyopathy and metabolic diseases linked to both disrupted iron homeostasis and impaired MLC function^1,2,33,34^.

## Methods

### Antibodies and plasmids

The following antibodies were used: PPS39 polyclonal antibody (Thermo Fisher Scientific; PA5-21104. ABclonal; A13082), TOMM22 monoclonal antibody (Proteintech; 66562-1-lg. Abcam; ab179826), SFXN1 polyclonal antibody (Merck; HPA019543. Proteintech; 68024-1-Ig), Alpha Tubulin Recombinant monoclonal antibody (Proteintech; 80762-1-R), HA Tag Recombinant monoclonal antibody (Proteintech; 81290-1-RR), mCherry Polyclonal antibody (Proteintech; 26765-1-AP), GFP polyclonal antibody (Abcam; ab290), STAR GREEN, goat anti-mouse IgG (Abberior; STGREEN-1001-500UG), STAR ORANGE, goat anti-rabbit IgG (Abberior; STORANGE-1002-500UG), Goat Anti-Rabbit IgG H&L (HRP) (Abcam; ab205718), and Goat Anti-Mouse IgG H&L (HRP) (Abcam; ab205719). Sequences of newly generated plasmids are provided in Supplementary Data 1.

### Cell culture

U2OS cells (BeNa Culture Collection) were cultured in McCoy’s 5A medium (Thermo Fisher Scientific) supplemented with 10% (v/v) fetal bovine serum (FBS; Thermo Fisher Scientific). HEK293T cells (BeNa Culture Collection) were cultured in Dulbecco’s modified Eagle’s medium (DMEM; Thermo Fisher Scientific) supplemented with 10% FBS. All cells were maintained at 37℃ in a humidified environment containing 5% CO_2_.

### Bioinformatics Analysis and Protein Screening

GO-CC Gene Ontology cellular component (GO CC) enrichment analysis was performed using Metascape (http://metascape.org). Candidate proteins were filtered from IP-mass spectrometry (MS) and ion transport datasets (https://www.gseamsigdb.org/gsea/msigdb/human/geneset/GOBP_IRON_ION_TRANSPORT) according to protein score, sequence coverage, and log_10_-transformed iBAQ abundance.

### Cell transfection and gene manipulation

Cells were transfected with fluorescent-tagged plasmids, overexpression constructs, target-specific small interfering RNA (siRNA), or short hairpin RNA (shRNA) using Lipofectamine™ 3000 or RNAiMax (Invitrogen) following the manufacturer’s protocol. Stable gene knockdown cell lines were generated following puromycin selection. Cells were harvested for Western blotting at 72 h after transfection, while parallel cultures were processed for imaging at 24 h post-transfection.

### Western Blot (WB) and Co-Immunoprecipitation (Co-IP)

For WB, cell lysates were prepared in RIPA buffer (Beyotime, P0013C) supplemented with protease inhibitor. Protein concentrations were determined using the BCA assay, and 30 μg of protein per lane was separated by SDS-PAGE and transferred to PPDF membranes (Merck Millipore). Membranes were blocked with 5% BSA in TBST and incubated with primary antibodies followed by HRP-conjugated secondary antibodies. Signals were detected using ECL substrate (Thermo Fisher Scientific, WP20005). For Co-IP, cells were lysed in ice-cold IP lysis buffer, and cleared lysates were incubated with Anti-GFP affinity beads 4FF (Smart-Lifesciences; SA070005), mCherry Nanoab Agarose Beads (NuoyiBio; CNA-25-500), or Anti-HA Magnetic Beads (MCE; HY-K0201) at 4 °C. After washing, bound proteins were eluted, separated by SDS-PAGE, and detected by WB to confirm protein-protein interactions.

### Fluorescent probe labeling in live cells

Cells were grown on glass-bottomed culture dishes (Biofil) before live-cell labeling. The following commercially available fluorescent probes were used according to the manufacturer’s protocols: RhoNox-M (Lumiprobe; 3317-50ug), Mito-FerroGreen (Dojindo; M489), FerroOrange (Dojindo; F374), LysoBrite red (AAT Bioquest; 22645), PK Mito deep red (Genvivo; PKMDR-1), Lysotracker Green (Thermo Fisher Scientific; L7526), and MitoTracker Green (Thermo Fisher Scientific; M7514). Morph-Atto488, and Atto647N labeling were performed as previously described^35^.

### Intracellular Fe(II) measurements

Intracellular Fe(II) levels were quantified using an Iron Assay Kit (Beyotime, S1066S) in accordance with the manufacturer’s instructions. Absorbance was measured at 593 nm using a microplate reader.

### Drug treatments

For drug treatments, live cells were labeled with FerroOrange for cytosol labile Fe(II), Mito-FerroGreen or RhoNox-M for mitochondrial or lysosomal Fe(II), respectively. LysoBrite Red and PK Mito Deep Red were used to label lysosomes and mitochondria, respectively. Following probe loading, cells were transferred to a custom serine-depleted medium supplemented with L-serine at the indicated concentrations (0.01, 0.1, 1, 10, or 100 mM), or treated with ammonium iron(II) sulfate hexahydrate (FAS) (100 μM; Sigma-Aldrich, 203505) or deferoxamine (DFO) (100 μM; Sigma-Aldrich, D9533) for 30 min, and subsequently subjected to live-cell imaging.

### Immunofluorescence (IF) and proximity ligation assay (PLA)

Cells were seeded into glass-bottomed culture dishes before fixed with 4% (m/v) paraformaldehyde (PFA; Electron Microscopy Sciences) for 13 min at 37℃ or pre-cooled methanol for 7 min at −20°C. PFA fixed cells were permeabilized with 0.2% (v/v) Triton X-100 (Sigma-Aldrich) for 5 min at room temperature (RT). Fixed cells were blocked in 5% goat serum (Thermo Fisher Scientific) for 1 h at RT. For IF staining, samples were incubated with primary antibodies overnight at 4 °C, followed by fluorophore-conjugated secondary antibodies for 1 h at room temperature. PLA was using the Duolink PLA kit (Merck; DUO92102-1KT) following the manufacturer’s protocol. Cells were stained with MitoTracker Green (Thermo Fisher Scientific) after PLA protocol to outline mitochondrial network.

### Microscopy and image acquisition

Confocal imaging was performed on a Nikon C2 laser-scanning confocal microscope equipped with a 60× oil-immersion objective (NA 1.49). Wide-field and structured illumination microscopy (SIM) imaging were carried out on a HIS-SIM system (CSR Biotech) fitted with a 100× oil-immersion objective (NA 1.50). For quantitative analysis of single-probe fluorescence intensity, wide-field microscopy was used to avoid intensity bias introduced by SIM reconstruction algorithms. For measuring direct Fe(II) transfer in dual-color SIM imaging (Fig. 1, Fig. 3, Extended Data Fig. 7), raw frames were acquired with an exposure time of 30 ms per frame without inter-frame intervals. Long-term SIM imaging of lysosome-mitochondria contacts (MLCs) (Fig. 2, Extended Data Fig. 4) was performed with a 3-s time interval and a 20 ms exposure time per frame.

### Image analysis

Fluorescence intensities of ion-sensitive probes were quantified using ImageJ software (National Institutes of Health, NIH). MLC events were quantified with defined criteria for counts, length and duration as indicated. Statistical comparisons were performed using paired/unpaired two-tailed Student’s t-test for two-group comparisons and one-way/two-way ANOPA followed by Tukey’s post-hoc test for multiple-group comparisons. Data are presented as mean ± s.e.m. unless otherwise stated. All statistical analyses were conducted with n ≥ 9 per condition from at least three independent biological replicates, as detailed in figure legends. Statistical graphs were generated using Microsoft Excel, and schematic illustrations were created with BioGDP.com^36^.

## Data Availability

All data that support the findings of this study are included in the manuscript or are available from the authors upon reasonable request.

## Code Availability

The custom-written Fiji-macro script for MLC quantification of the first frames is available at https://github.com/AllenHoo-sea/Fiji-Lysosome-Mitochondria-Contact, and the custom written MATLAB workflow for MLC total events, MLC duration, and lysosomes quantification in long-term dual-color SIM imaging is available at https://github.com/iolee963/MLC-Time-and-Length-Analysis.

## Acknowledgements

We thank Wenwen Gong and Weiyun Sun for experimental and technical assistance. We thank the Optical Bioimaging Core Facility of WNLO-HUST (Wuhan National Laboratory for Optoelectronics-Huazhong University of Science and Technology), the Research Core Facilities for Life Science of HUST, and Advanced Biomedical Imaging Facility-WNLO for the support in data acquisition.

## Fundings

This work was supported by the National Natural Science Foundation of China (62125504, 32671819, 62675288, 32271428), National Key R&D Program of China (2022YFC3401100), Zhejiang Provincial Natural Science Foundation of China (LZ25F050008), Leading Innovative and Entrepreneur Team Introduction Program of Zhejiang (2024R01001)

## Author contributions

Y.H., C.K., and Y.-H.Z. conceived the project. Y.H. and X.H. designed experiments, analyzed data, and wrote the manuscript with input from all authors. Y.H. performed cell culture, immunofluorescence, PLA, and confocal imaging. X.H. performed Western Blot (WB), wide-field imaging, and structured illumination microscopy (SIM) experiments. X.P. performed SIM imaging experiments. X.H. and R.Q. developed the algorithms for image analysis. All authors contributed to the data analysis.

## Competing interests

The authors declare no competing interests.

## Additional information

**Supplementary information** Supplementary Information is available for this paper. **Correspondence and requests for materials** should be addressed to Y.H., C.K., or Y.-H.Z.

## Extended Data Figure

**Extended Data Fig. 1.**
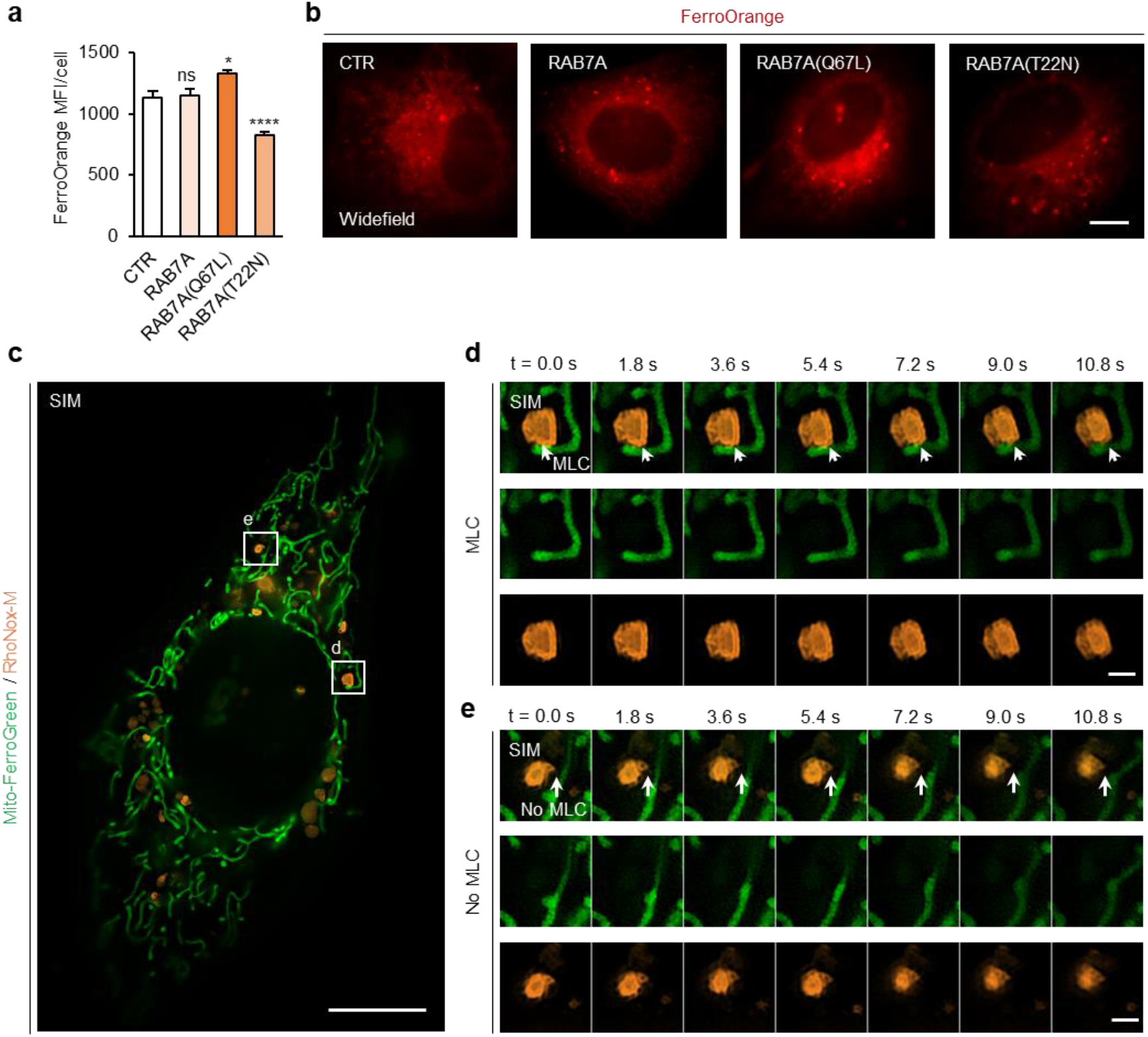
Cytosolic labile iron pool (LIP) measurements and MLC iron dynamic time-lapse imaging. **a**, **b**, Quantification (**a**; from left to right: n =93, 105, 105, 105 cells) and representative widefield images (**b**) of cytosolic labile iron pool (LIP; FerroOrange) under the indicated conditions. Scale bars, 10 μm. **c–e**, Representative dual-color SIM images of Mito-FerroGreen and RhoNox-M co-staining. Whole-cell overview (**c**). Scale bar, 10 μm. Magnified time-lapse snapshots of MLC (**d**) and no-MLC (**e**) regions from **c**. Scale bars, 1 μm. Statistical analysis was performed using one-way ANOPA with Tukey’s multiple comparisons test. *P < 0.05, **P < 0.01, ***P < 0.001, ****P < 0.0001; ns, not significant.

**Extended Data Fig. 2.**
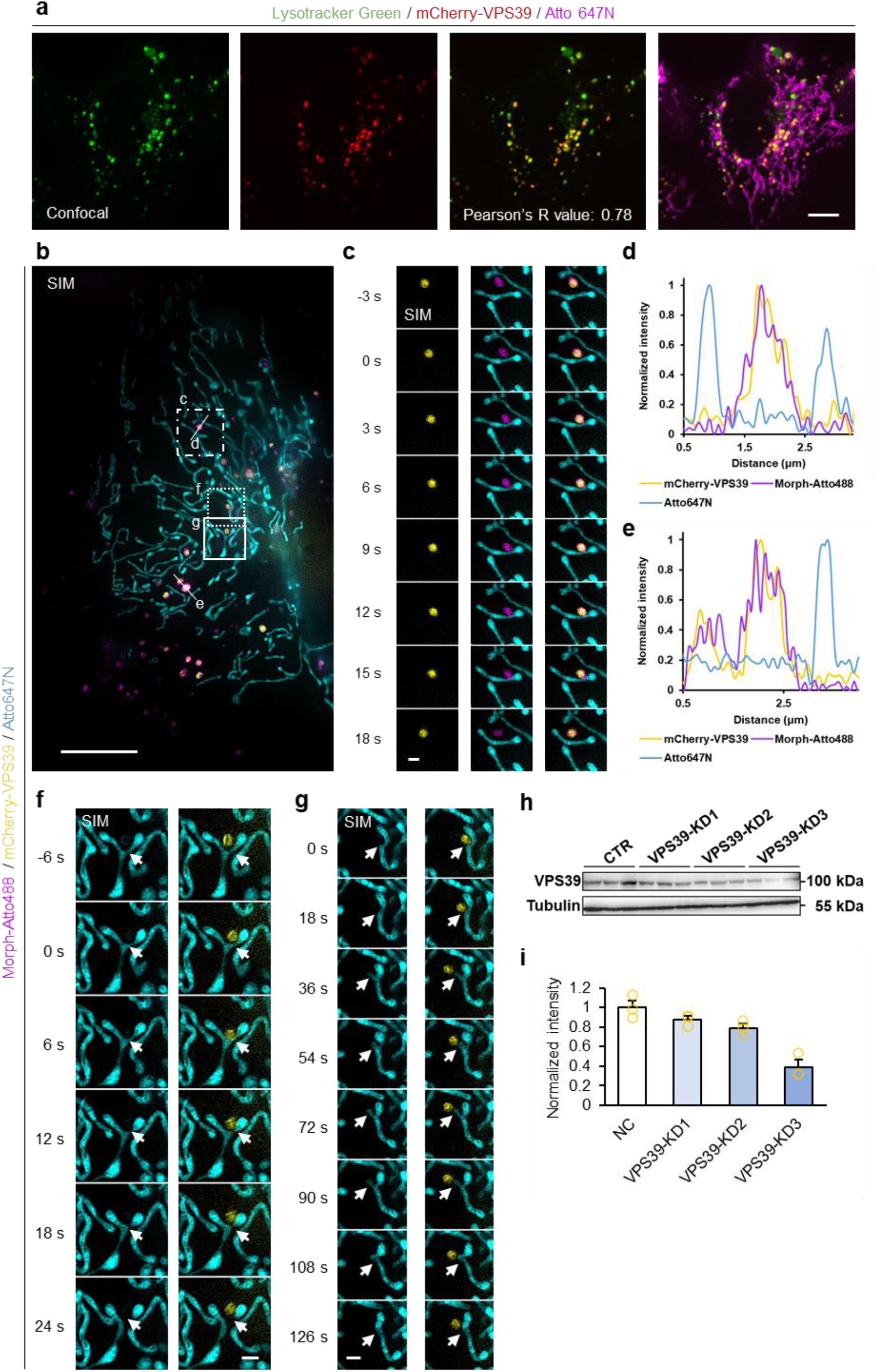
Intracellular localization, dynamic characteristics, and expression profile of VPS39. **a**, Three-color confocal images of live cells stained with LysoTracker Green (green), mCherry-PPS39 (red), and Atto 647N (magenta). Colocalization between PPS39 and lysosomes was observed (Pearson’s coefficient = 0.78). Scale bar, 10 μm. **b–g**, Three-color time-lapse SIM of live cells stained with Morph-Atto488 (magenta), mCherry-PPS39 (yellow), and Atto 647N (cyan). First-frame of the time-lapse series (**b**; scale bar, 10 μm). Magnified views of boxed regions in **b** from the time-lapse SIM series (**c**, **f**, **g**; scale bars, 1 μm). Multicolor intensity profiles (**d**, **e**) along the lines indicated in **b**. **h**, **i**, Western blot (WB) images (h) and corresponding quantitative analysis (i) of PPS39 expression levels.

**Extended Data Fig. 3.**
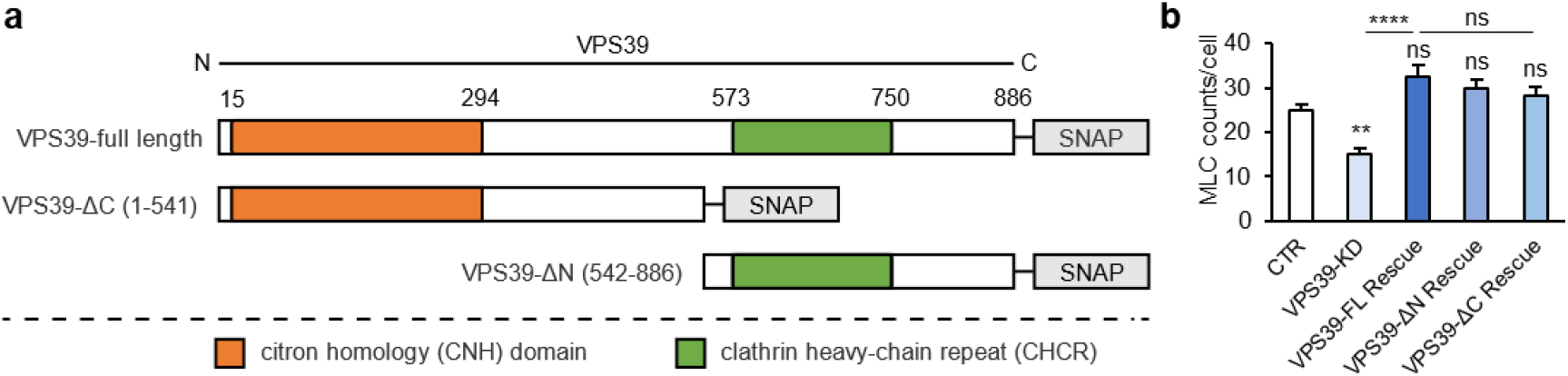
Domain mapping of VPS39 for the regulation of mitochondrial-lysosome contact (MLC) formation. **a**, Schematic diagram of truncated SNAP-tagged fragments and the predicted domain organization of PPS39 protein. Domain boundaries were annotated according to conserved domain information from UniProt and Pfam databases. **b**, Quantification of MLC abundance following rescue expression of full-length and truncated PPS39 constructs (from left to right: n = 102, 98, 102, 102, 103).

**Extended Data Fig. 4.**
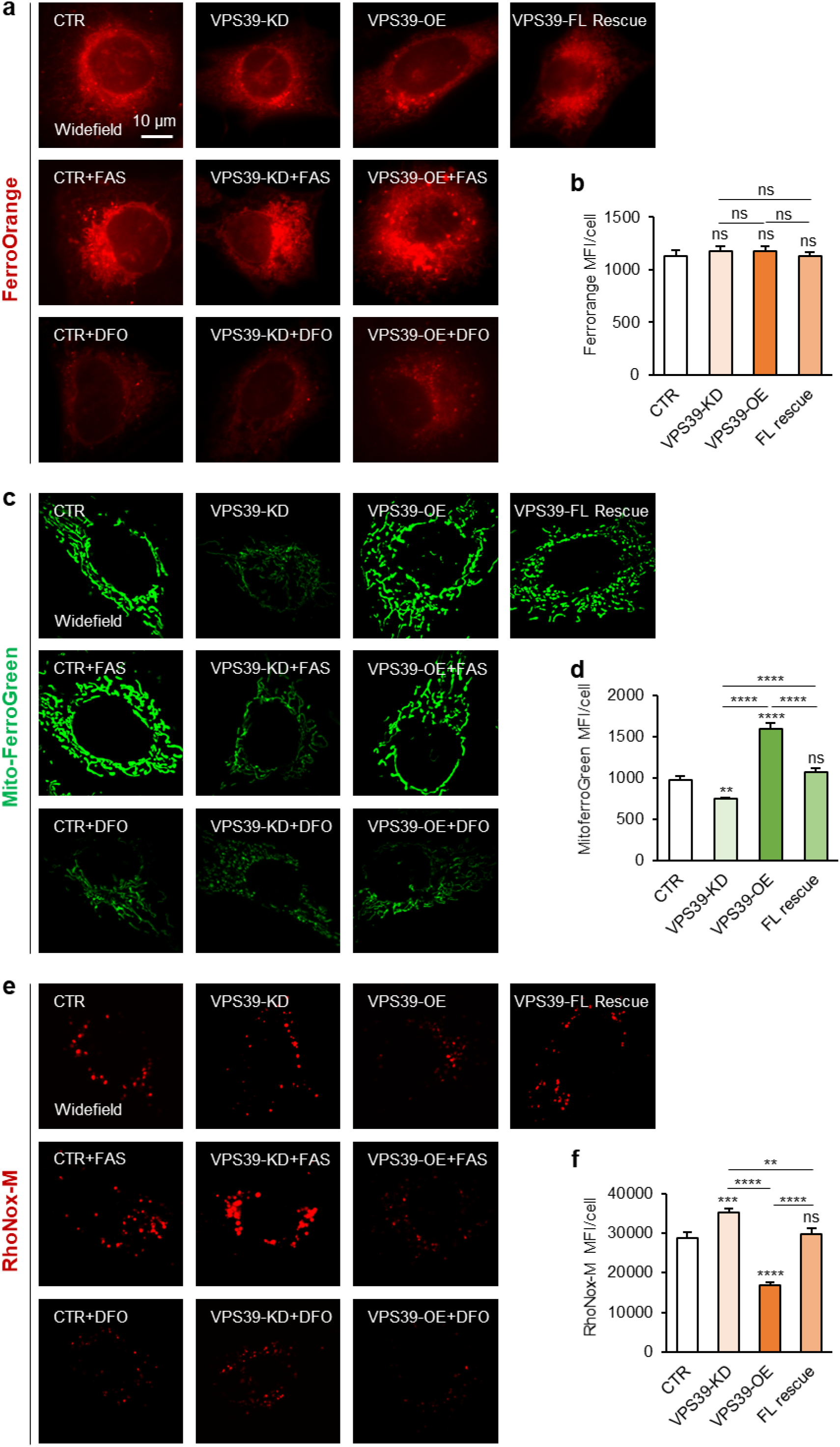
Fe(II) levels in mitochondria, lysosomes, and the labile iron pool (LIP) under different protein A levels with FAS or DFO treatment. **a**, **c**, **e**, Representative images of LIP (**a**; FerroOrange), lysosomal iron (**c**; RhoNox-M), and mitochondrial iron (**e**; Mito-FerroGreen) under the indicated conditions. Scale bar, 10 μm. **b**, **d**, **f**, Quantification of LIP (**b**; from left to right: n = 115, 109, 105, 105 cells), mitochondrial iron (**f**; from left to right: n = 110, 123, 105, 96 cells), and lysosomal iron (**d**; from left to right: n = 102, 110, 106, 103 cells) under the indicated conditions. Statistical analysis was performed using one-way ANOPA with Tukey’s multiple comparisons test. *P < 0.05, **P < 0.01, ***P < 0.001, ****P < 0.0001; ns, not significant.

**Extended Data Fig. 5.**
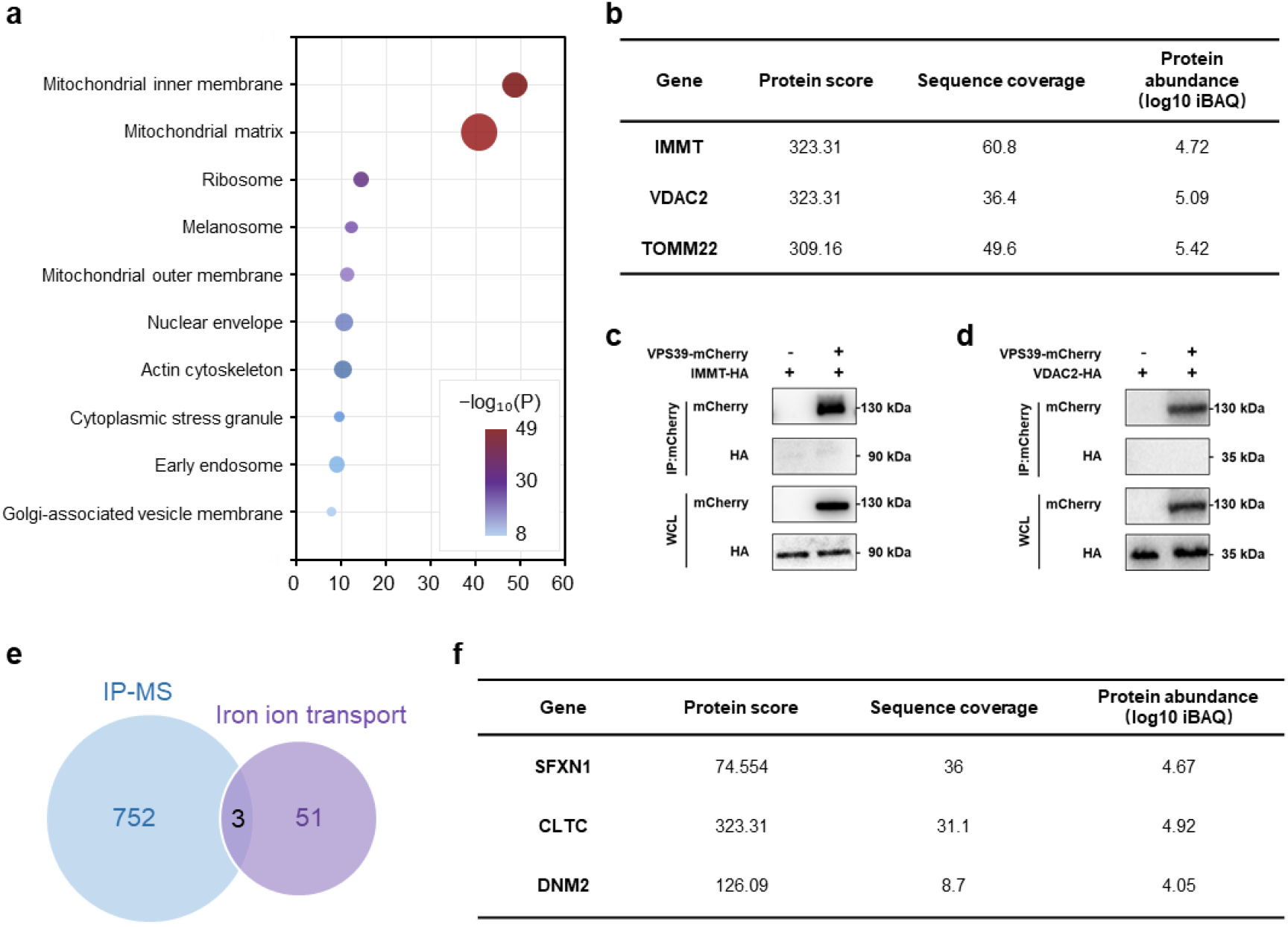
Spatial association among VPS39, TOMM22 and SFXN1. **a**, Gene Ontology Cellular Component (GO-CC) analysis of PPS39 interactors. Bubble size corresponds to the number of enriched genes (counts); color intensity indicates −log_10_-transformed P-values. **b**, Mass spectrometry identification of three candidate PPS39-interacting proteins on the outer mitochondrial membrane (OMM). **c**, **d**, Co-immunoprecipitation (co-IP) of PPS39 with IMMT (**c**) and PDAC2 (**d**). **e**, **f**, Penn diagram (**f**) and table (**g**) showing the overlap between the PPS39 interactome mass spectrometry library and iron transporter datasets.

**Extended Data Fig. 6.**
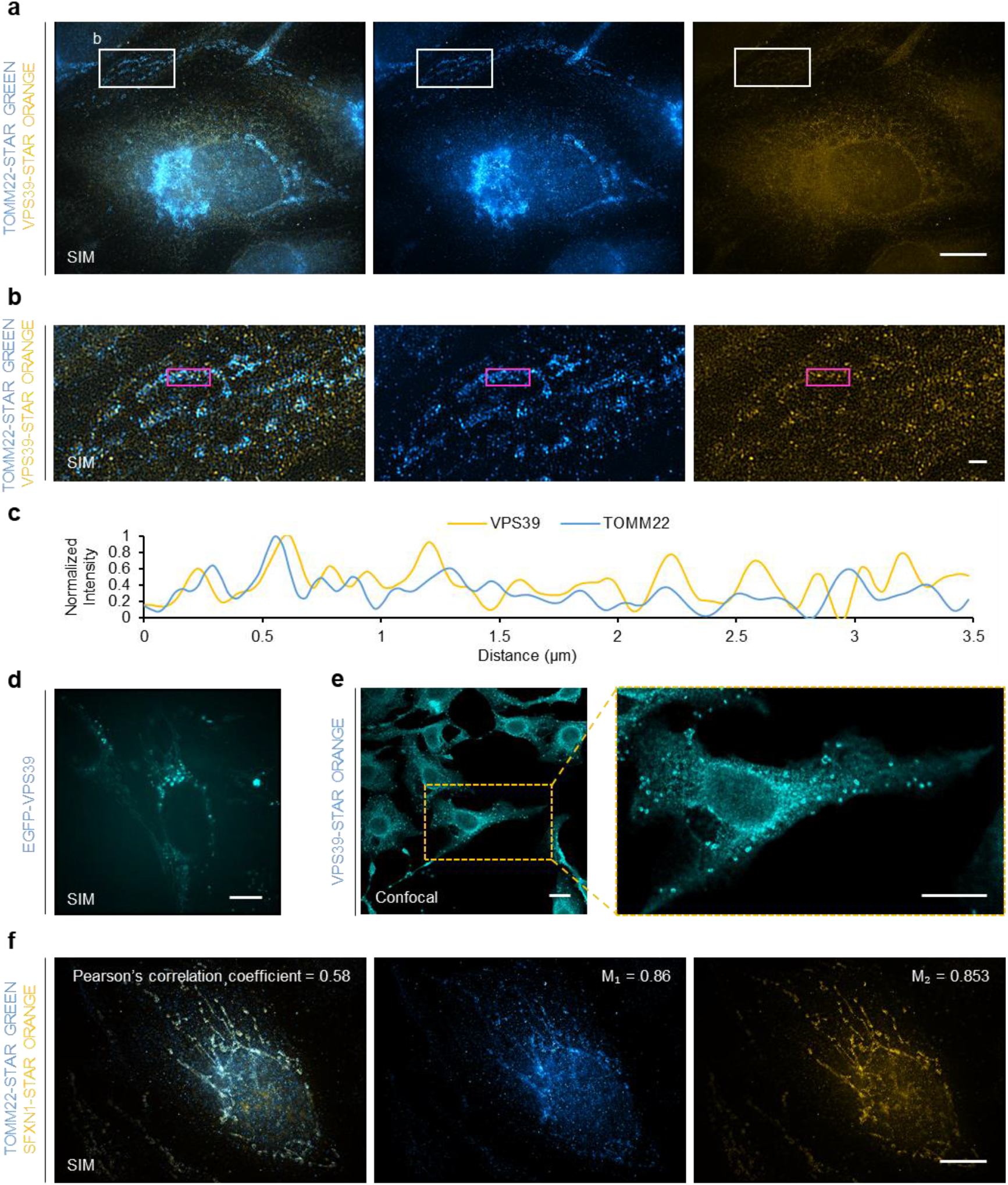
Dual-color SIM analysis of VPS39 and SFXN1 colocalization with TOMM22. **a–c**, Dual-color SIM immunofluorescence imaging of PPS39 and TOMM22 (**a**; fixed in ice-cold methanol; scale bar, 10 μm. **b**; magnified view of the white-boxed region in a, scale bar, 1 μm). Fluorescence intensity line profile (**c**) corresponding to the magenta boxed region indicated in (**b**). **d**, **e**, Representative images of EGFP-PPS39 in live-cell SIM (**d**; scale bar, 10 μm) and PPS39 immunostaining in confocal microscopy (**e**; scale bars, 20 μm), showing punctate distribution and mitochondrial-like morphology. **f,** Representative SIM images and quantitative colocalization analysis between SFXN1 and TOMM22 (fixed in ice-cold methanol; scale bar, 10 μm).

**Extended Data Fig. 7.**
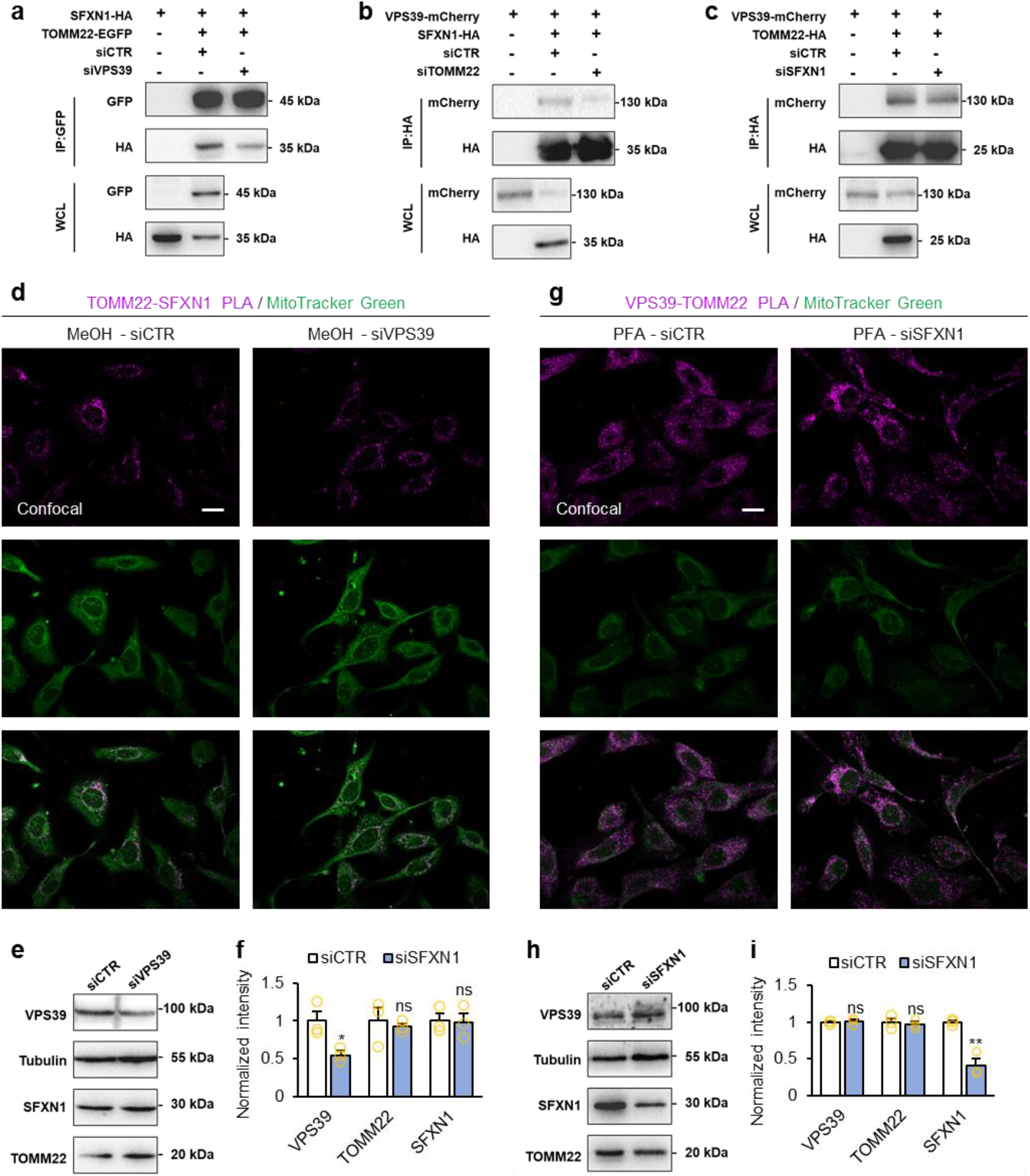
Modulation of pairwise interactions among TOMM22, SFXN1 and VPS39 by siRNA silencing of the third component. **a–c,** Co-immunoprecipitation (co-IP) assays assessing the interactions between the indicated protein pairs following siRNA-mediated silencing of PPS39 (**a**), TOMM22 (**b**) or SFXN1 (**c**). **d**, **g**, Representative proximity ligation assay (PLA) images showing PPS39-TOMM22 and TOMM22-SFXN1 interactions. Methanol-fixed cells under siCTR and siPPS39 conditions were used for TOMM22-SFXN1 detection (**d**), whereas PFA-fixed cells under siCTR and siSFXN1 conditions were used for PPS39-TOMM22 detection (**g**). Scale bars, 20 μm. **e–i**, Western-blot images and corresponding quantification of PPS39, TOMM22 and SFXN1 following knockdown of PPS39 (**e, f**), and SFXN1 (**h, i**). Data represent three independent biological replicates. Statistical analysis was performed using two-tailed unpaired Student’s t-test. *P < 0.05, **P < 0.01, ***P < 0.001, ****P < 0.0001; ns, not significant.

**Extended Data Fig. 8.**
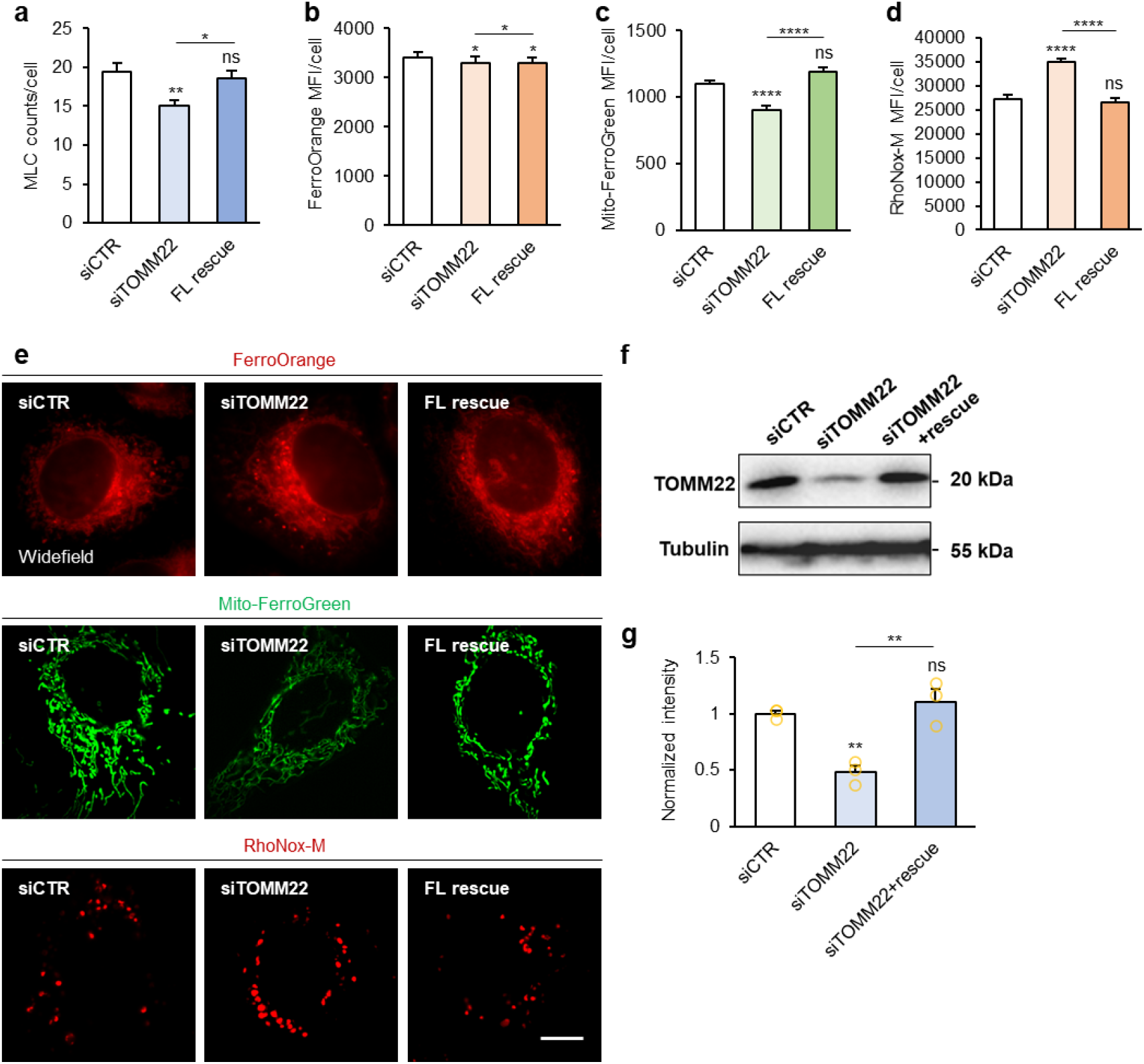
TOMM22 acts as the essential bridging factor for the ternary complex assembly and Fe(II) transfer. **a–e**, Quantification of MLC counts (**a**; from left to right: n = 95, 104, 104 cells) and iron levels in the LIP (**b**; from left to right: n = 105, 105, 105 cells), mitochondria (**c**; from left to right: n = 105, 105, 102 cells), and lysosomes (**d**; from left to right: n = 105, 105, 105 cells) and representative widefield images (**e**) under the indicated conditions. Scale bar, 10 μm. **f**, **g**, Western blot (WB) images (**f**) and corresponding quantitative analysis (**g**) of TOMM22 expression upon siRNA-mediated knockdown of TOMM22 and other indicated conditions. Statistical analysis was performed using one-way ANOPA with Tukey’s multiple comparisons test. *P < 0.05, **P < 0.01, ***P < 0.001, ****P < 0.0001; ns, not significant.

**Extended Data Fig. 9.**
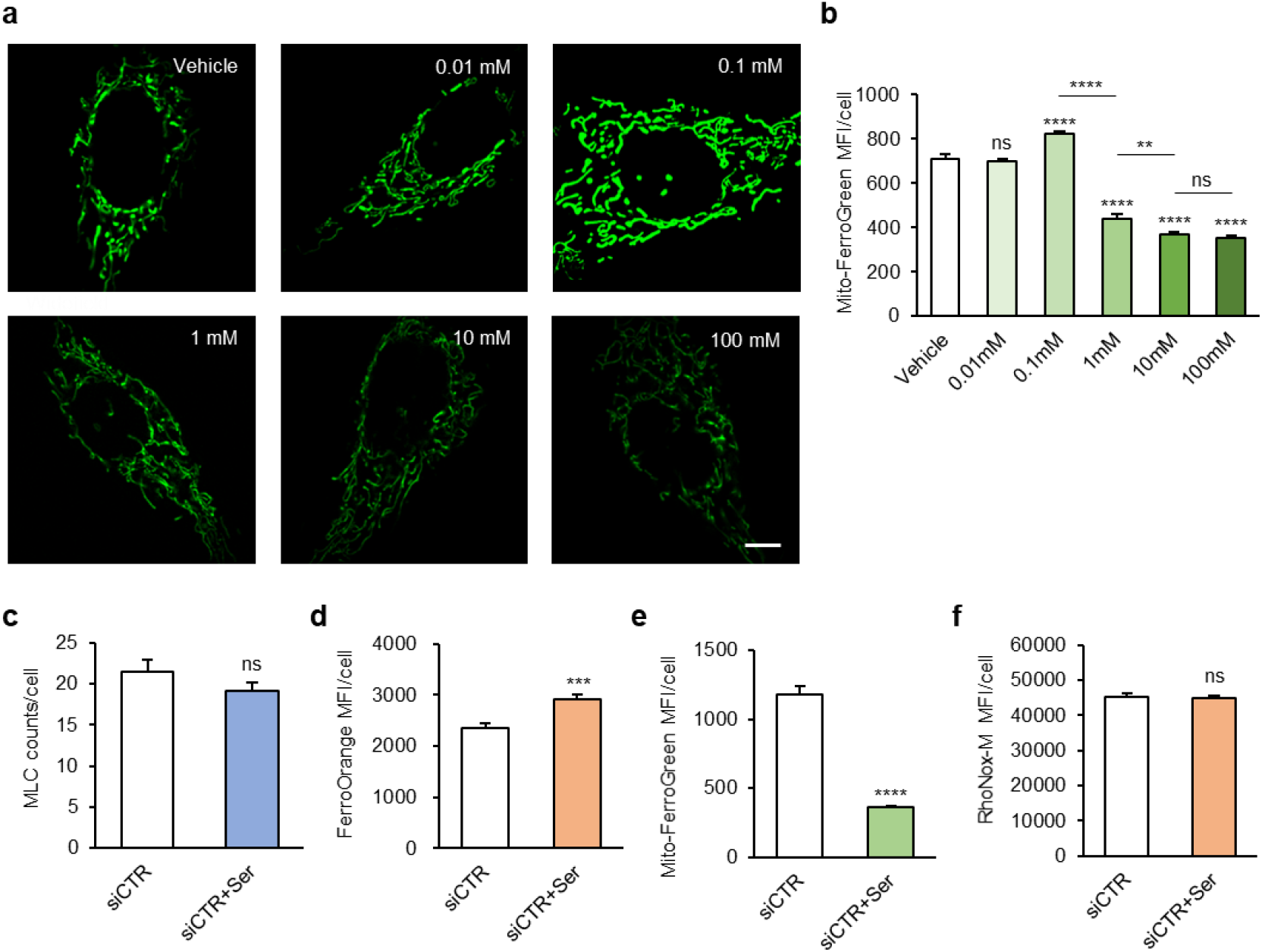
MLC abundance and subcellular labile iron levels upon L-serine treatment. **a**, Representative fluorescence images stained with Mito-FerroGreen after treatment with vehicle or L-serine at indicated concentrations (0.01, 0.1, 1, 10, and 100 mM) for 0.5 h. Scale bar = 10 μm. **b**, Quantification of relative Mito-FerroGreen fluorescence intensity from the images shown in (**a**). From left to right: n = 102, 104, 103, 104, 105, 105 cells. Statistical analysis was performed using one-way ANOPA with Tukey’s multiple comparisons test. **c–d**, Quantification of MLC counts (**a**; from left to right: n = 105 and 104 cells), ferrous iron levels in the LIP (**b**; from left to right: n = 99 and 101 cells), mitochondria (**c**; from left to right: n = 109 and 105 cells), and lysosomes (**d**; from left to right: n = 105 and 105 cells) under indicated conditions. Statistical analysis was performed using using two-tailed unpaired Student’s t-test. *P < 0.05, **P < 0.01, ***P < 0.001, ****P < 0.0001; ns, not significant.

**Extended Data Fig. 10.**
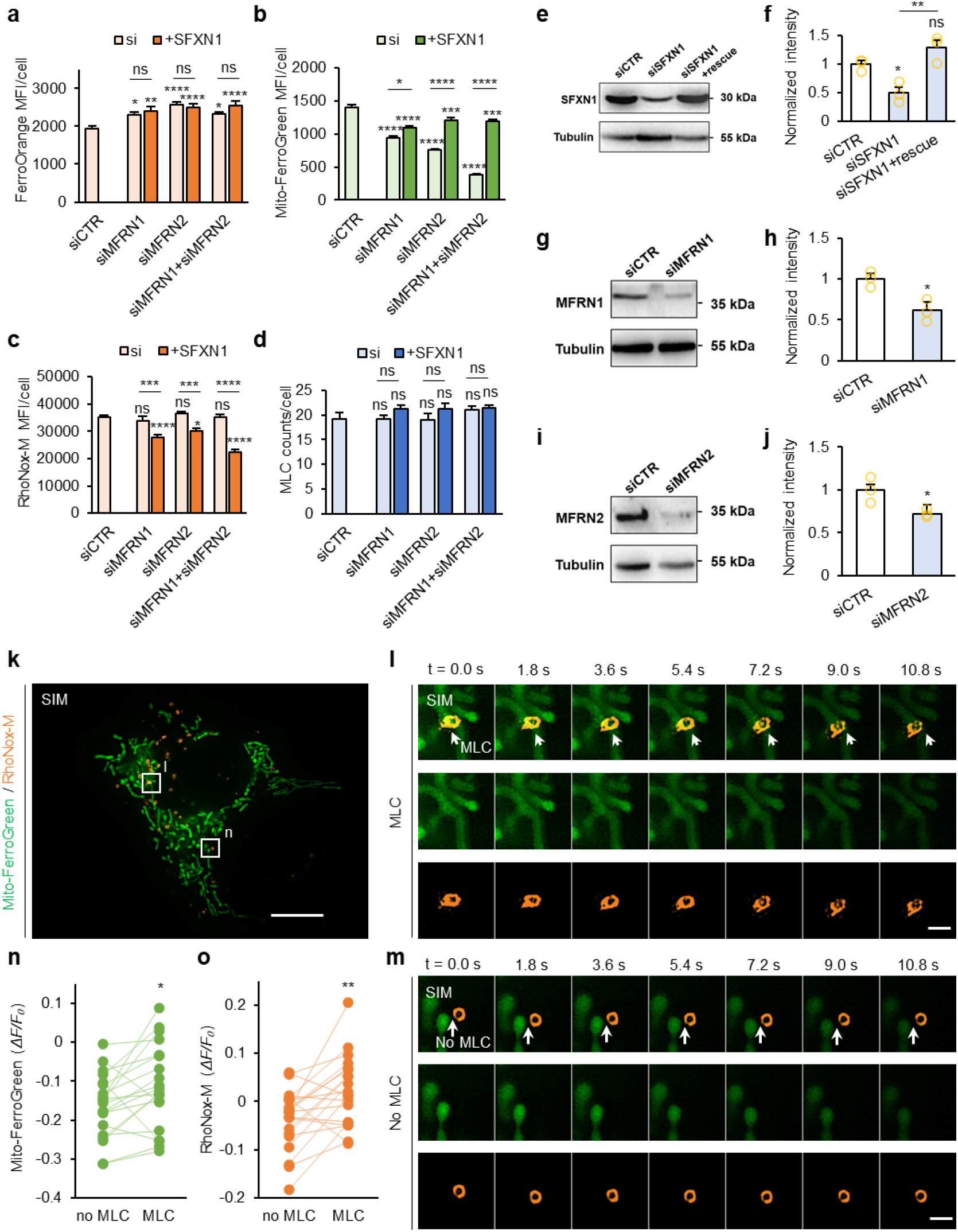
SFXN1 depletion impairs lysosome-to-mitochondria Fe(II) transfer. **a–d**, Quantification of ferrous iron levels in the LIP (**c**; from left to right: n = 100, 102, 103, 102, 102, 103, 102 cells), mitochondria (**d**; from left to right: n = 101, 97, 105, 105, 105, 105, 105 cells), lysosomes (**e**; from left to right: n = 105, 105, 105, 105, 105, 105, 105, 105, 105 cells), and MLC counts (**f**; from left to right: n = 101, 88, 86, 99, 92, 86, 87 cells) upon siRNA-mediated knockdown of MFRN1, MFRN2, or both. **e–j**, Western blot images and corresponding quantitative analysis of SFXN1 (**e, f**), MFRN1 (**g**, **h**), and MFRN2 (**i**, **j**) expression upon indicated conditions. **k–o**, Representative dual-color SIM images of Mito-FerroGreen and RhoNox-M co-staining in siSFXN cells. Whole-cell overview (**k**); scale bar, 10 μm. Magnified time-lapse snapshots of MLC (**l**) and no-MLC (**m**) regions from **k**; scale bars, 1 μm. Normalized fluorescence change (*ΔF/F₀*) values of Mito-FerroGreen (**n**) and RhoNox-M (**o**) signals were compared between MLC sites and paired no-MLC regions (n= 20). Fluorescence signals were photobleaching-corrected using intensity from no-MLC areas. Statistical analysis was performed using one-way ANOPA with Tukey’s multiple comparisons test (**a–d**, **f**), two-tailed unpaired Student’s t-test (**h**, **j**), or two-tailed paired Student’s t-test (**n**, **o**). *P < 0.05, **P < 0.01, ***P < 0.001, ****P < 0.0001; ns, not significant.

## Notes

### Competing Interest Statement

The authors have declared no competing interest.

