## Supplemental Information for "A direct, MFRN-independent Fe(II) transfer pathway at mitochondria-lysosome contacts"

### Supplementary Information

**Supplementary Note 1. Mechanistic analysis of VPS39-mediated lysosome-mitochondria Fe(II) trafficking under drug treatment.**

Under untreated conditions, cytosolic LIP showed no obvious differences among groups (Fig. 2g–i). VPS39 knockdown reduced mitochondrial Fe(II) content while elevating lysosomal Fe(II) level, and VPS39 overexpression exerted the opposite effects. These results indicate that VPS39 regulates Fe(II) transport between lysosomes and mitochondria.

**VPS39-dependent Fe(II) distribution upon ferrous ammonium sulfate (FAS) supplementation**

FAS treatment elevated cytosolic LIP Fe(II) in all groups, independent of VPS39 expression.

**CTR+FAS vs. CTR untreated:** In control cells, mitochondrial Fe(II) was markedly higher in FAS-treated samples than in untreated ones, while lysosomal Fe(II) exhibited only a slight increase via indirect trafficking pathways. This indicates that the elevated cytoplasmic LIP is rapidly delivered to mitochondria, which is consistent with the known function of FAS[1].

**KD+FAS vs. KD untreated:** In FAS-treated VPS39 knockdown cells, lysosomal Fe(II) was elevated compared with the untreated KD group. This suggests that VPS39 depletion does not prevent excess cytoplasmic Fe(II) from accumulating in lysosomes. Meanwhile, mitochondrial Fe(II) levels were comparable across groups. This implies that upon mitochondria-lysosome contact (MLC) disruption, mitochondria fail to take up extra Fe(II), even when cytoplasmic and lysosomal Fe(II) pools are abundant.

**KD+FAS vs. CTR+FAS:** Cytosolic LIP exhibited no statistical difference between control and KD groups upon FAS stimulation, demonstrating VPS39 deficiency fails to alter overall cytoplasmic labile Fe(II) abundance. Compared with FAS-treated control cells, KD group presented accumulated lysosomal Fe(II) and decreased mitochondrial Fe(II), further verifying that the absence of MLC blocks Fe(II) transport from lysosomes to mitochondria and eventually causes Fe(II) retention in lysosomes.

**OE+FAS vs. OE untreated:** No obvious changes in lysosomal and mitochondrial Fe(II) were detected in VPS39-overexpressed cells regardless of FAS administration, indicating Fe(II) transport between the two organelles reaches saturated equilibrium upon excessive VPS39 expression, and redundant Fe(II) is reserved in cytoplasmic LIP.

**OE+FAS vs. CTR+FAS:** Compared with FAS-treated control group, OE group had higher mitochondrial Fe(II) and lower lysosomal Fe(II), supporting that abundant VPS39 facilitates Fe(II) export from lysosomes to mitochondria. And mitochondrial Fe(II) saturation may correspondingly diminishes Fe(II) uptake from cytoplasm, leading to the accumulation of surplus Fe(II) in LIP.

Collectively, under FAS treatment, lysosome-mitochondria Fe(II) trafficking depends mainly on VPS39 levels. On the other hand, Fe(II) flux from cytosolic LIP to mitochondria remains unresponsive to VPS39 manipulation, even in the presence of FAS.

**VPS39-dependent Fe(II) distribution under Deferoxamine (DFO) chelation**

Upon DFO treatment, lysosomal Fe(II) levels decreased relative to the untreated group.

**CTR+DFO vs. CTR untreated:** Cytosolic LIP showed no marked changes under DFO treatment, suggesting that DFO may indirectly and gradually reduce the intracellular labile Fe(II) pool. Mitochondrial Fe(II) declined slightly, likely as a secondary consequence of reduced lysosomal Fe(II).

**CTR+DFO vs. CTR untreated & KD+DFO vs. KD untreated:** After DFO treatment, mitochondrial Fe(II) decreased mildly in normal control cells, whereas it remained nearly unchanged in VPS39 KD cells. This suggests that the reduction in mitochondrial Fe(II) upon DFO treatment originates largely from lysosomal Fe(II) pool in control cells, and this regulatory effect is diminished when MLC function is impaired in KD cells. Notably, the DFO-mediated reduction in lysosomal Fe(II) did not affect the residual mitochondrial Fe(II) in VPS39 KD cells. These findings suggest that, under severe Fe(II) deficiency, cells prioritize mitochondrial Fe(II) supply through cytosolic LIP-to-mitochondria trafficking. Cytosolic LIP remained unchanged in DFO-treated control cells but decreased in KD cells, which supports the compensatory Fe(II) supply from LIP to mitochondria. Lysosomal Fe(II) was higher in DFO-treated KD group than control group, suggesting that blocked MLC retains Fe(II) that should be transported to mitochondria.

**OE+ DFO vs. OE untreated vs. CTR+ DFO:** Under DFO-induced lysosomal Fe(II) depletion, VPS39 OE strengthened Fe(II) export toward mitochondria and further consumed lysosomal Fe(II), accompanied by marked reduction of cytoplasmic LIP. Restricted total lysosomal Fe(II) content lowered mitochondrial Fe(II) level in OE cells relative to untreated OE group, while mitochondrial Fe(II) was comparable between OE and control groups after DFO treatment. It indicates cells sustain maximal mitochondrial Fe(II) supply even with limited lysosomal Fe(II) reserve.

Collectively, DFO intervention led to comparable mitochondrial Fe(II) levels in control, knockdown and overexpression cells. These observations suggest that when total cellular Fe(II) is limited, cells employ multiple pathways involving cytosolic LIP and lysosomes to retain basal mitochondrial Fe(II) for physiological homeostasis.

**Supplementary Note 2. Detailed screening process of candidate proteins involved in MLC-mediated lysosome-to-mitochondria Fe(II) transport**

We first performed Gene Ontology Cellular Component (GO-CC) enrichment on VPS39 protein-associated interactors (Extended Data Fig. 3a). The analysis revealed strong enrichment in mitochondrial compartments. This bioinformatic evidence suggests functional crosstalk between lysosomes and mitochondria, implying that other mitochondrial interacting proteins may cooperate with VPS39 to modulate MLC-associated biological processes. We then selected three top-ranked candidate proteins located on the outer mitochondrial membrane (OMM) for further validation via Co-IP (Extended Data Fig. 3b). The results confirmed that only TOMM22 interacted with VPS39, whereas the other two candidates showed no detectable interaction (Fig. 3a and Extended Data Fig. 3c,d). We further performed Co-IP assays and confirmed that TOMM22 interacts with the C-terminal region of VPS39, but not the N-terminal region.

Given that TOMM22 functions as a protein channel rather than an ion transporter, we hypothesize that the VPS39-TOMM22 complex primarily mediates MLC formation via molecular bridging, while additional proteins are required to execute actual ion transport. Accordingly, we intersected the interactome of VPS39 and the dataset of iron ion transport proteins and got three candidate proteins (Extended Data Fig. 3e,f). After strict screening based on subcellular localization, protein function and literature evidence, we focused on SFXN1[23], an inner mitochondrial membrane (IMM) protein with conserved iron ion translocation activity. Subsequent biochemical experiments confirmed that SFXN1 interacts with both VPS39 and TOMM22, supporting the existence of a V-T-S ternary interaction (Fig. 3b,c and Extended Data Fig. 3g–i). This stepwise screening and logical deduction help bridge the mechanistic gap between MLC structural assembly and ion transport function, and offer a preliminary framework and working hypothesis for subsequent functional characterization of the three proteins.

**Supplementary Note 3. Detailed discussion about immunofluorescence (IF) results.**

Fixation regimens were optimized based on antigen characteristics of each target protein. Our results showed that methanol fixation is required to preserve SFXN1 immunoreactivity, as PFA crosslinking causes severe signal loss of SFXN1. PFA fixation generates superior staining quality for the outer mitochondrial membrane protein TOMM22. Accordingly, PFA fixation was used for dual-color staining of VPS39 and TOMM22, while methanol fixation was applied for dual-color co-staining of SFXN1 and TOMM22to preserve detectable SFXN1 signals.

Although endogenous and GFP-tagged VPS39 exhibited signals tracing mitochondrial networks (Fig. 4g and Extended Data Fig. 6a–e), this faint mitochondrial VPS39 pool could not be visualized in SIM acquisitions using mCherry-VPS39 (Fig. 2a,b and Extended Data Fig. 2). EGFP generally has a higher quantum yield and undergoes faster chromophore maturation relative to mCherry[4]. These biochemical properties likely enable EGFP-fused VPS39 to reflect this low-abundance mitochondrial population during SIM imaging. By comparison, mCherry tends to undergo photobleaching more readily during serial SIM scanning, and its comparatively weaker detection sensitivity may obscure the faint mitochondrial VPS39 signal above background levels.

The ring-shaped, lysosome-like staining pattern of VPS39 was not consistently reproduced across all immunostaining batches. Our data suggest that increasing the primary antibody concentration to 1:100 enhances the sensitivity of this ring-like staining, yet this concentration concurrently resulted in elevated background signals in proximity ligation assay (PLA). Therefore, the antibody concentration was titrated to 1:200 to achieve an optimal signal-to-noise ratio in dual-color staining. Meanwhile, the variable lysosomal localization of VPS39 may reflect its sensitivity to cellular physiological states, consistent with its role in HOPS complex-mediated membrane fusion[5].

**Supplementary Table 1. Statistical results of FerroOrange via two-way ANOVA with Tukey's multiple comparisons test.**

| Tukey's multiple comparisons test | Summary | Adjusted P Value |
|---|---|---|
| CTR:Untreated vs CTR:FAS | **** | <0.0001 |
| CTR:Untreated vs CTR:DFO | ns | 0.0616 |
| CTR:Untreated vs VPS39-KD:Untreated | ns | 0.9994 |
| CTR:Untreated vs VPS39-KD:FAS | ** | 0.0036 |
| CTR:Untreated vs VPS39-KD:DFO | ** | 0.0012 |
| CTR:Untreated vs VPS39-OE:Untreated | ns | 0.9988 |
| CTR:Untreated vs VPS39-OE:FAS | **** | <0.0001 |
| CTR:Untreated vs VPS39-OE:DFO | **** | <0.0001 |
| CTR:FAS vs CTR:DFO | **** | <0.0001 |
| CTR:FAS vs VPS39-KD:Untreated | **** | <0.0001 |
| CTR:FAS vs VPS39-KD:FAS | ns | 0.5516 |
| CTR:FAS vs VPS39-KD:DFO | **** | <0.0001 |
| CTR:FAS vs VPS39-OE:Untreated | **** | <0.0001 |
| CTR:FAS vs VPS39-OE:FAS | ** | 0.0023 |
| CTR:FAS vs VPS39-OE:DFO | **** | <0.0001 |
| CTR:DFO vs VPS39-KD:Untreated | ** | 0.0092 |
| CTR:DFO vs VPS39-KD:FAS | **** | <0.0001 |
| CTR:DFO vs VPS39-KD:DFO | ns | 0.9749 |
| CTR:DFO vs VPS39-OE:Untreated | ** | 0.0082 |
| CTR:DFO vs VPS39-OE:FAS | **** | <0.0001 |
| CTR:DFO vs VPS39-OE:DFO | **** | <0.0001 |
| VPS39-KD:Untreated vs VPS39-KD:FAS | * | 0.0378 |
| VPS39-KD:Untreated vs VPS39-KD:DFO | **** | <0.0001 |
| VPS39-KD:Untreated vs VPS39-OE:Untreated | ns | >0.9999 |
| VPS39-KD:Untreated vs VPS39-OE:FAS | **** | <0.0001 |
| VPS39-KD:Untreated vs VPS39-OE:DFO | **** | <0.0001 |
| VPS39-KD:FAS vs VPS39-KD:DFO | **** | <0.0001 |
| VPS39-KD:FAS vs VPS39-OE:Untreated | ns | 0.0504 |
| VPS39-KD:FAS vs VPS39-OE:FAS | **** | <0.0001 |
| VPS39-KD:FAS vs VPS39-OE:DFO | **** | <0.0001 |
| VPS39-KD:DFO vs VPS39-OE:Untreated | **** | <0.0001 |
| VPS39-KD:DFO vs VPS39-OE:FAS | **** | <0.0001 |
| VPS39-KD:DFO vs VPS39-OE:DFO | **** | <0.0001 |
| VPS39-OE:Untreated vs VPS39-OE:FAS | **** | <0.0001 |
| VPS39-OE:Untreated vs VPS39-OE:DFO | **** | <0.0001 |
| VPS39-OE:FAS vs VPS39-OE:DFO | **** | <0.0001 |

**Supplementary Table 2. Statistical results of Mito-FerroGreen via two-way ANOVA with Tukey’s multiple comparisons test.**

| Tukey's multiple comparisons test | Summary | Adjusted P Value |
|---|---|---|
| CTR:Untreated vs. CTR:FAS | **** | <0.0001 |
| CTR:Untreated vs. CTR:DFO | ** | 0.0076 |
| CTR:Untreated vs. VPS39-KD:Untreated | ** | 0.0034 |
| CTR:Untreated vs. VPS39-KD:FAS | ** | 0.0019 |
| CTR:Untreated vs. VPS39-KD:DFO | **** | <0.0001 |
| CTR:Untreated vs. VPS39-OE:Untreated | **** | <0.0001 |
| CTR:Untreated vs. VPS39-OE:FAS | **** | <0.0001 |
| CTR:Untreated vs. VPS39-OE:DFO | *** | 0.0009 |
| CTR:FAS vs. CTR:DFO | **** | <0.0001 |
| CTR:FAS vs. VPS39-KD:Untreated | **** | <0.0001 |
| CTR:FAS vs. VPS39-KD:FAS | **** | <0.0001 |
| CTR:FAS vs. VPS39-KD:DFO | **** | <0.0001 |
| CTR:FAS vs. VPS39-OE:Untreated | **** | <0.0001 |
| CTR:FAS vs. VPS39-OE:FAS | **** | <0.0001 |
| CTR:FAS vs. VPS39-OE:DFO | **** | <0.0001 |
| CTR:DFO vs. VPS39-KD:Untreated | ns | >0.9999 |
| CTR:DFO vs. VPS39-KD:FAS | ns | >0.9999 |
| CTR:DFO vs. VPS39-KD:DFO | ns | 0.9229 |
| CTR:DFO vs. VPS39-OE:Untreated | **** | <0.0001 |
| CTR:DFO vs. VPS39-OE:FAS | **** | <0.0001 |
| CTR:DFO vs. VPS39-OE:DFO | ns | 0.9997 |
| VPS39-KD:Untreated vs. VPS39-KD:FAS | ns | >0.9999 |
| VPS39-KD:Untreated vs. VPS39-KD:DFO | ns | 0.9708 |
| VPS39-KD:Untreated vs. VPS39-OE:Untreated | **** | <0.0001 |
| VPS39-KD:Untreated vs. VPS39-OE:FAS | **** | <0.0001 |
| VPS39-KD:Untreated vs. VPS39-OE:DFO | ns | >0.9999 |
| VPS39-KD:FAS vs. VPS39-KD:DFO | ns | 0.9909 |
| VPS39-KD:FAS vs. VPS39-OE:Untreated | **** | <0.0001 |
| VPS39-KD:FAS vs. VPS39-OE:FAS | **** | <0.0001 |
| VPS39-KD:FAS vs. VPS39-OE:DFO | ns | >0.9999 |
| VPS39-KD:DFO vs. VPS39-OE:Untreated | **** | <0.0001 |
| VPS39-KD:DFO vs. VPS39-OE:FAS | **** | <0.0001 |
| VPS39-KD:DFO vs. VPS39-OE:DFO | ns | 0.9983 |
| VPS39-OE:Untreated vs. VPS39-OE:FAS | ns | >0.9999 |
| VPS39-OE:Untreated vs. VPS39-OE:DFO | **** | <0.0001 |
| VPS39-OE:FAS vs. VPS39-OE:DFO | **** | <0.0001 |

**Supplementary Table 3. Statistical results of RhoNox-M via two-way ANOVA with Tukey's multiple comparisons test.**

| Tukey's multiple comparisons test | Summary | Adjusted P Value |
|---|---|---|
| CTR:Untreated vs CTR:FAS | * | 0.0219 |
| CTR:Untreated vs CTR:DFO | **** | <0.0001 |
| CTR:Untreated vs VPS39-KD:Untreated | *** | 0.0001 |
| CTR:Untreated vs VPS39-KD:FAS | **** | <0.0001 |
| CTR:Untreated vs VPS39-KD:DFO | **** | <0.0001 |
| CTR:Untreated vs VPS39-OE:Untreated | **** | <0.0001 |
| CTR:Untreated vs VPS39-OE:FAS | **** | <0.0001 |
| CTR:Untreated vs VPS39-OE:DFO | **** | <0.0001 |
| CTR:FAS vs CTR:DFO | **** | <0.0001 |
| CTR:FAS vs VPS39-KD:Untreated | ns | 0.9481 |
| CTR:FAS vs VPS39-KD:FAS | **** | <0.0001 |
| CTR:FAS vs VPS39-KD:DFO | **** | <0.0001 |
| CTR:FAS vs VPS39-OE:Untreated | **** | <0.0001 |
| CTR:FAS vs VPS39-OE:FAS | **** | <0.0001 |
| CTR:FAS vs VPS39-OE:DFO | **** | <0.0001 |
| CTR:DFO vs VPS39-KD:Untreated | **** | <0.0001 |
| CTR:DFO vs VPS39-KD:FAS | **** | <0.0001 |
| CTR:DFO vs VPS39-KD:DFO | **** | <0.0001 |
| CTR:DFO vs VPS39-OE:Untreated | **** | <0.0001 |
| CTR:DFO vs VPS39-OE:FAS | **** | <0.0001 |
| CTR:DFO vs VPS39-OE:DFO | ns | 0.2335 |
| VPS39-KD:Untreated vs VPS39-KD:FAS | **** | <0.0001 |
| VPS39-KD:Untreated vs VPS39-KD:DFO | **** | <0.0001 |
| VPS39-KD:Untreated vs VPS39-OE:Untreated | **** | <0.0001 |
| VPS39-KD:Untreated vs VPS39-OE:FAS | **** | <0.0001 |
| VPS39-KD:Untreated vs VPS39-OE:DFO | **** | <0.0001 |
| VPS39-KD:FAS vs VPS39-KD:DFO | **** | <0.0001 |
| VPS39-KD:FAS vs VPS39-OE:Untreated | **** | <0.0001 |
| VPS39-KD:FAS vs VPS39-OE:FAS | **** | <0.0001 |
| VPS39-KD:FAS vs VPS39-OE:DFO | **** | <0.0001 |
| VPS39-KD:DFO vs VPS39-OE:Untreated | ns | >0.9999 |
| VPS39-KD:DFO vs VPS39-OE:FAS | ns | 0.9998 |
| VPS39-KD:DFO vs VPS39-OE:DFO | **** | <0.0001 |
| VPS39-OE:Untreated vs VPS39-OE:FAS | ns | 0.9998 |
| VPS39-OE:Untreated vs VPS39-OE:DFO | **** | <0.0001 |
| VPS39-OE:FAS vs VPS39-OE:DFO | **** | <0.0001 |

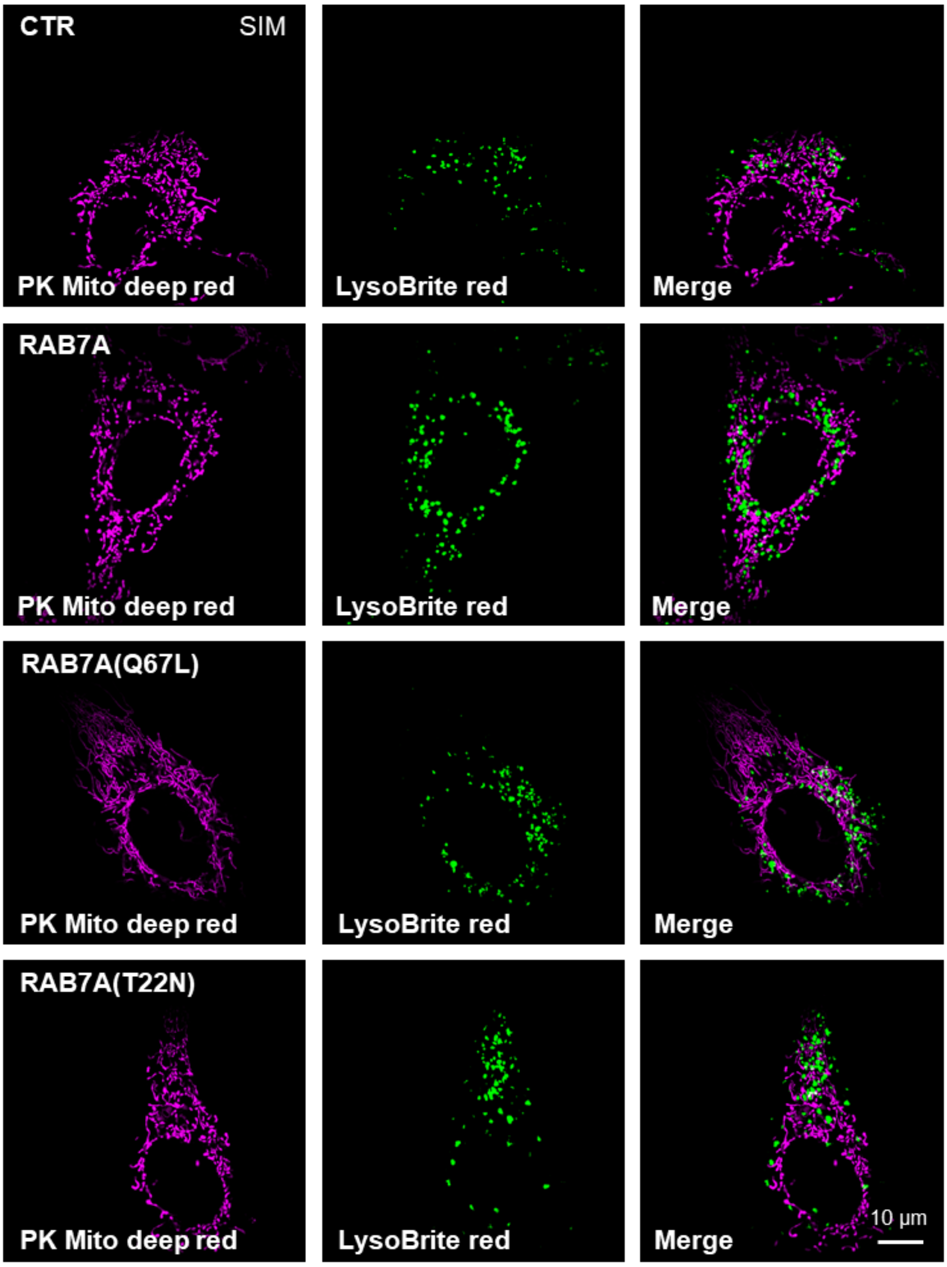

**Supplementary Figure 1. Structured illumination microscopy (SIM) imaging of mitochondria-lysosome contacts (MLCs).**

Representative SIM images of MLCs under the indicated conditions.

Fig3a

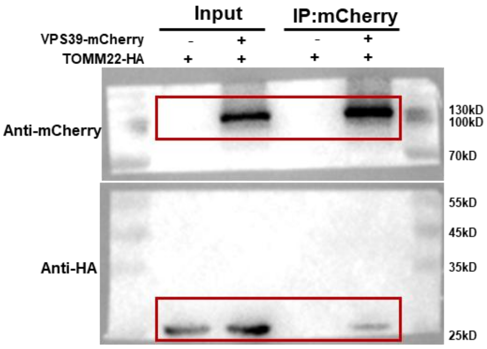

Fig3b

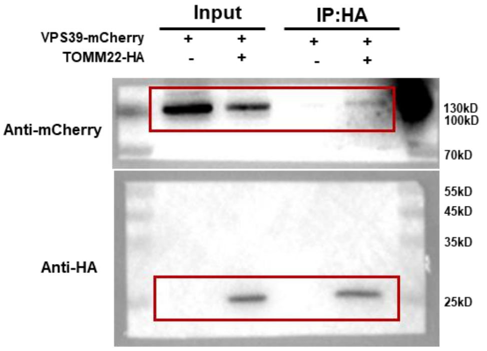

Fig3c

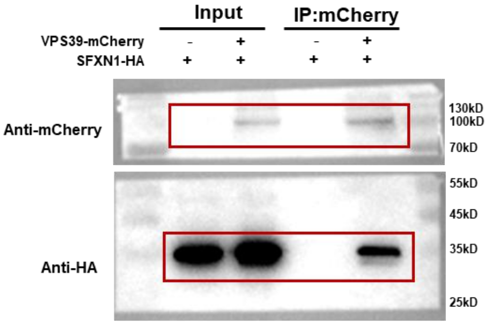

Fig3d

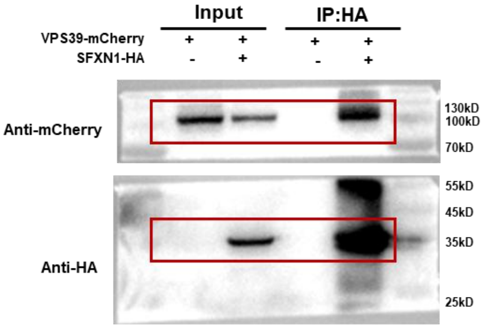

Fig3e

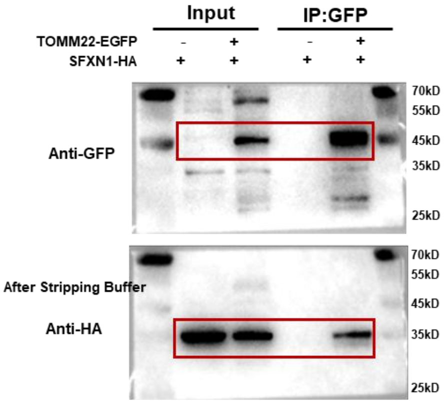

Fig3f

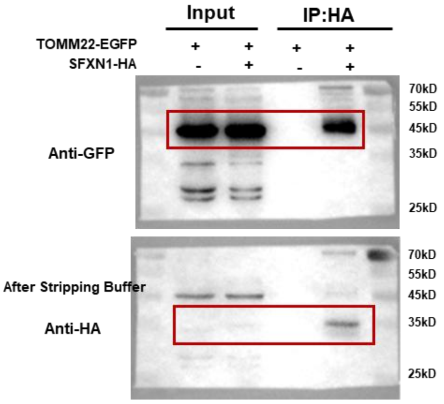

#### Extended Data Fig2h

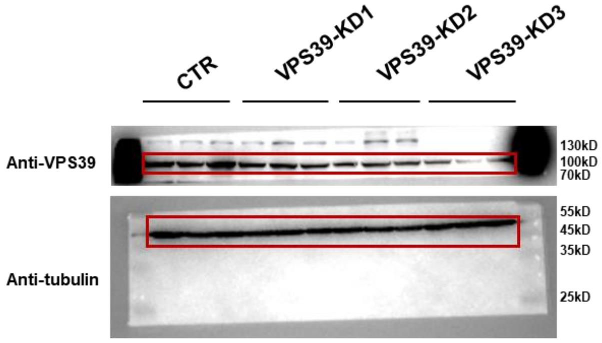

#### Extended Data Fig5c

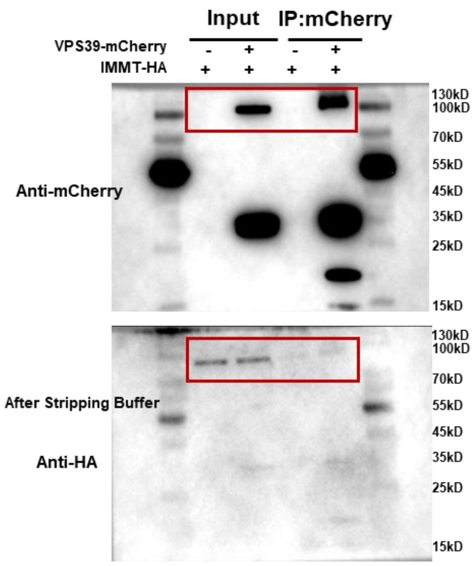

#### Extended Data Fig5d

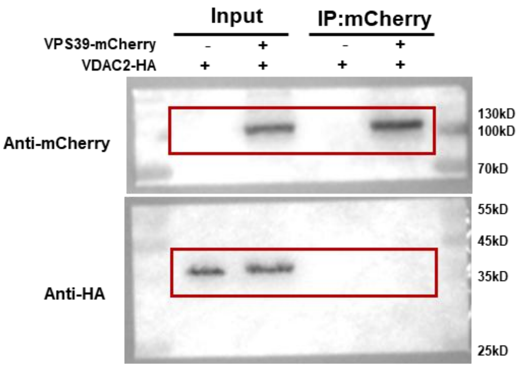

##### Extended Data Fig7a

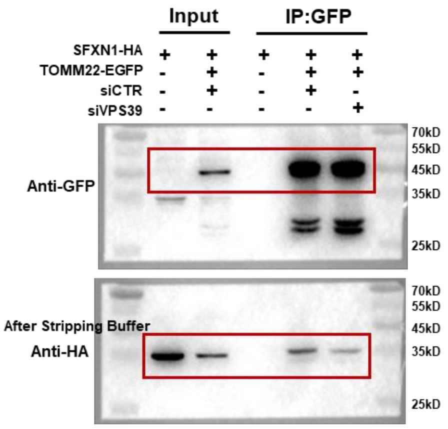

##### Extended Data Fig7b

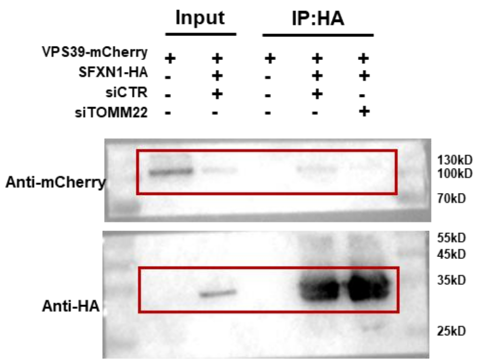

##### Extended Data Fig7c

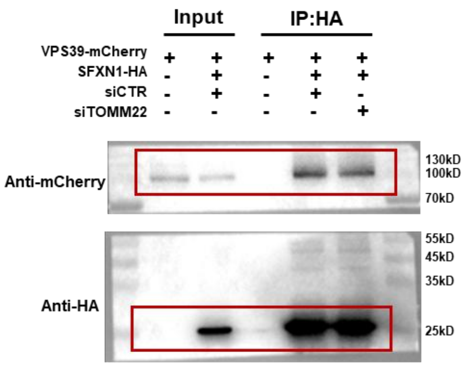

##### Extended Data Fig7e

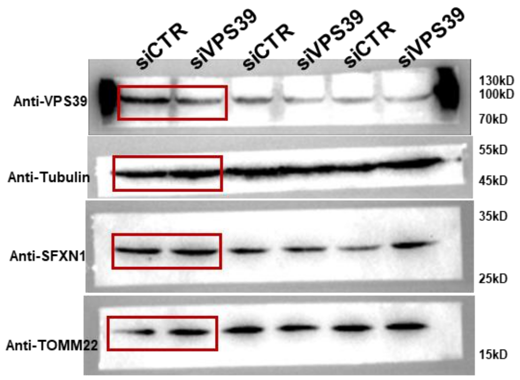

##### Extended Data Fig7h

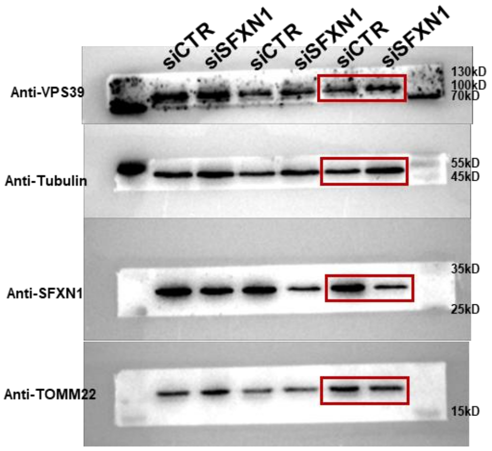

#### Extended Data Fig8f

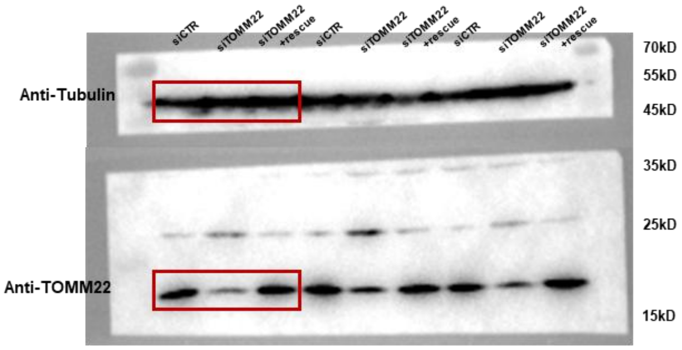

#### Extended Data Fig10e

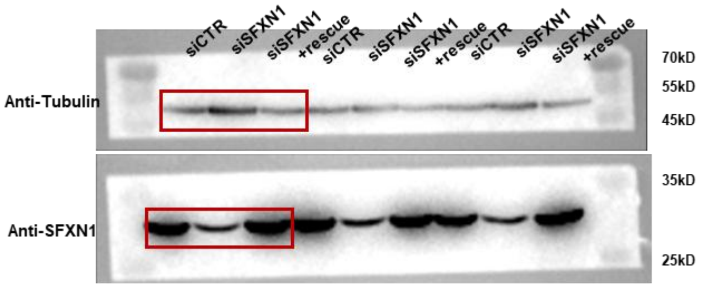

#### Extended Data Fig10g

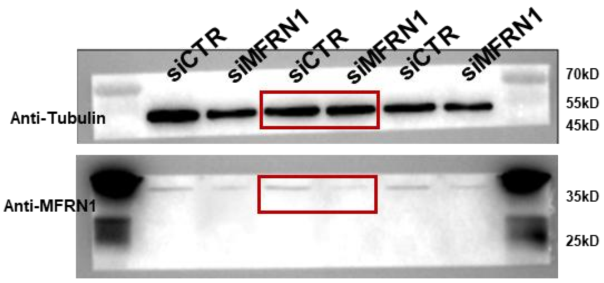

#### Extended Data Fig10i

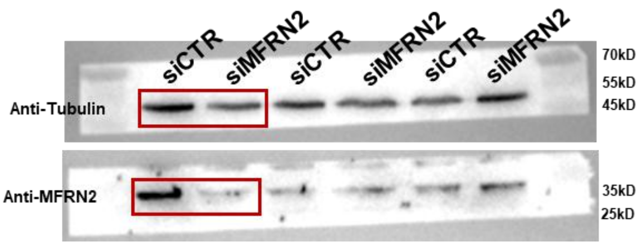

**Supplementary Figure 2. Uncropped western blot images.**

**Supplementary Data 1: Sequences of plasmids generated and used in this study.**

To construct the mCherry-VPS39 recombinant plasmid, the human VPS39 fragment was amplified from cDNA and inserted into a mammalian expression vector. Subsequently, EGFP-VPS39 was generated by replacing the mCherry coding sequence in the mCherry-VPS39 construct with an EGFP sequence. This mCherry-VPS39 plasmid also served as the template to construct the lentiviral vector pLV2-CMV-VPS39-IRES-SNAP and its truncation mutants pLV2-CMV-VPS39-(1-541aa)-IRES-SNAP and pLV2-CMV-VPS39-(542-875aa)-IRES-SNAP. Additional expression plasmids used in this study included pCMV-SNAP-RAB7A-Neo (G99585), pCMV-SNAP-Rab7A-Q67L-Neo (G99587), pCMV-SNAP-2×HA-Rab7A-T22N-Neo (G99586), pCMV-IMMT-3×HA-Neo (P89970), pCMV-SFXN1-3×HA-Neo (G107086), pCMV-TOMM22-3×HA-Neo (G95491), pCMV-TOMM22-EGFP-Neo (G108885), and pCMV-VDAC2-3×HA-Neo (G95490), all of which were obtained from Wuhan Miaoling Bioscience and Technology (Wuhan, China).

siRNAs (Sangon Biotech) and shRNAs (WZ Bioscience Inc.) were used for gene silencing. All sequences are shown in the 5'→3' direction. The target sequences are listed below:

siVPS39: GGCUAUUCCUUAUCUGGAA

siTOMM22: AUCAUGAAGGAAGUGGUCC

siSFXN1: GGGAACAGCUUACGUUUCUTT

siMitoferrin1: GCCATTTCTTGGTCTGTCTAT

siMitoferrin2: CCUGCUACGAAAAGUUAAA

VPS39-ShRNA1: CGTTGGTTTCAAGAGAGACTA

VPS39-ShRNA2: GGATGGAAAGGGGTCCATCAAH

VPS39-ShRNA3: CCACGTCTTGAAGGACACAAA
